# Diverse beta burst waveform motifs characterize movement-related cortical dynamics

**DOI:** 10.1101/2022.12.13.520225

**Authors:** Maciej J Szul, Sotirios Papadopoulos, Sanaz Alavizadeh, Sébastien Daligaut, Denis Schwartz, Jérémie Mattout, James J Bonaiuto

## Abstract

Classical analyses of induced, frequency-specific neural activity typically average bandlimited power over trials. More recently, it has become widely appreciated that in individual trials, beta band activity occurs as transient bursts rather than amplitude-modulated oscillations. Most studies of beta bursts treat them as unitary, and having a stereotyped waveform. However, we show there is a wide diversity of burst shapes. Using a biophysical model of burst generation, we demonstrate that waveform variability is predicted by variability in the synaptic drives that generate beta bursts. We then use a novel, adaptive burst detection algorithm to identify bursts from human MEG sensor data recorded during a joystick-based reaching task, and apply principal component analysis to burst waveforms to define a set of dimensions, or motifs, that best explain waveform variance. Finally, we show that bursts with a particular range of waveform motifs, ones not fully accounted for by the biophysical model, differentially contribute to movement-related beta dynamics. Sensorimotor beta bursts are therefore not homogeneous events and likely reflect distinct computational processes.

## Introduction

The spectral content and frequency band-specific modulations of neural signals have long served as fundamental building blocks for theories of how information is processed and propagated in the nervous system. Since early descriptions of such signals, activity in the beta frequency range (13 - 30 Hz) has been associated with movement preparation and performance (e.g. Jasper & Penfield, 1949; Murthy & Fetz, 1992; Pfurtscheller, 1981). In trial-averaged data, beta power gradually decreases before a movement, reaches a minimum during performance of the movement, and rapidly increases beyond baseline level following its completion (Alayrangues et al., 2019; Cassim et al., 2001; Cheyne, 2013; Donner et al., 2009; Erbil & Ungan, 2007; Haegens et al., 2011; Keinrath et al., 2006; Kilavik et al., 2013; Kilner et al., 2003; Leocani & Comi, 2006; McFarland et al., 2000; Meirovitch et al., 2015; K. J. Miller et al., 2010; Pfurtscheller et al., 1996; Pfurtscheller & Lopes da Silva, 1999; Pogosyan et al., 2009; Salenius & Hari, 2003; Tan et al., 2016; Tzagarakis et al., 2010, 2015). This signal has been implicated in a variety of motor processes including movement planning and preparation (Alayrangues et al., 2019; Bartolo & Merchant, 2015; Donner et al., 2009; Haegens et al., 2011; Heinrichs-Graham et al., 2016; Rhodes et al., 2018; Tzagarakis et al., 2015), inhibition (e.g. Cheyne, 2013; Jensen et al., 2005; Khanna & Carmena, 2017; Picazio et al., 2014; Pogosyan et al., 2009; van Wijk et al., 2009; Wessel & Aron, 2017; Y. Zhang et al., 2008), and learning (e.g. Boonstra et al., 2007; Fine et al., 2017; Houweling et al., 2008; Nakagawa et al., 2011; Pollok et al., 2014; Reuter et al., 2022; Tan et al., 2014), however the mechanism by which beta activity underlies this diverse range of suggested functions is not known.

Based on the temporal pattern of changes in trial-averaged beta power, this signal has been assumed to reflect sustained oscillatory activity in each trial, with movement-related amplitude modulations. However, it is becoming increasingly apparent that sensorimotor beta activity does not occur as sustained oscillations in single trials, but as transient burst events that occur with varying probability in time (Cagnan et al., 2019; Diesburg et al., 2021; Feingold et al., 2015; Haufler et al., 2022; Howe et al., 2011; Karvat et al., 2020; Kosciessa et al., 2020; Little et al., 2019; Lofredi et al., 2019; Sherman et al., 2016; Shin et al., 2017; Sporn et al., 2020; Torrecillos et al., 2018; Wessel, 2020). These variations in burst probability are reflected in the slow changes in average spectral power (Feingold et al., 2015; Howe et al., 2011; Little et al., 2019; Rayson et al., 2022; Sherman et al., 2016), but suggest that single trial beta activity is more dynamic than classically thought. The rate and timing of single-trial beta events are tightly linked to behavior (Diesburg et al., 2021; Echeverria-Altuna et al., 2021; Enz et al., 2021; Feingold et al., 2015; Haufler et al., 2022; Heideman et al., 2020; Howe et al., 2011; Karvat et al., 2020; Khawaldeh et al., 2020; Kosciessa et al., 2020; Law et al., 2022; Little et al., 2019; Sherman et al., 2016; Shin et al., 2017; Sporn et al., 2020; Torrecillos et al., 2018; Walsh et al., 2022; Wessel, 2020; West et al., 2022), and are more predictive of response time and accuracy than average beta amplitude (Enz et al., 2021; Little et al., 2019). This shift in perspective from trial-averaged beta power to transient burst events has thus allowed for more fine-grained analyses of their relationship to behavior, but the functional role of beta bursts themselves is still unclear.

Neural field potentials are commonly analyzed using Fourier or Hilbert transform-based approaches which assume that the activity is stationary, at least within a small time window, and sinusoidal (Cole & Voytek, 2017, 2019). Although such approaches are useful for isolating activity in different frequency channels, both of these assumptions are typically unmet by neural data. Therefore time-frequency (TF) based features of band-specific activity, whether based on trial-averaged power or bursts, cannot adequately capture nonsinusoidal dynamics in the temporal domain. However, recent work has shown that nonsinusoidal waveforms in the theta, alpha, and gamma bands are behaviorally (Marshall et al., 2022) and physiologically (Higgins et al., 2022; Quinn et al., 2021) informative, underscoring the importance of analyzing frequency-specific activity in the temporal domain without band pass filtering.

In sensorimotor cortex, beta bursts have a stereotypical wavelet-like shape in the temporal domain (Baker et al., 1997; Bonaiuto et al., 2021; Brady & Bardouille, 2022; Cagnan et al., 2019; Cole et al., 2017; Karvat et al., 2020; Kosciessa et al., 2020; Sherman et al., 2016) caused by temporally aligned synaptic drives to deep and superficial cortical layers (Bonaiuto et al., 2021; Sherman et al., 2016). The shape of this waveform determines the duration, peak amplitude, peak frequency, and frequency span of the burst in TF space. A large range of waveform shapes can potentially result in a burst with a peak frequency within the beta range when Fourier or Hilbert transform analyses are applied (Karvat et al., 2020; Sherman et al., 2016). Indeed, while the mean burst waveform shape appears highly conserved across studies, subjects, and species (Bonaiuto et al., 2021; Brady & Bardouille, 2022; Cagnan et al., 2019; Cole et al., 2017; Howe et al., 2011; Karvat et al., 2020; Kosciessa et al., 2020; Sherman et al., 2016), there is a large amount of variability in individual burst waveforms (Bonaiuto et al., 2021; Howe et al., 2011; Karvat et al., 2020; Kosciessa et al., 2020; Sherman et al., 2016). This waveform shape variability may translate to variability in function, potentially reconciling the various views of beta’s functional role by decomposing the signal into multiple distinct sources. However, the vast majority of previous work has treated beta bursts as being entirely homogeneous, and focused on burst rate, timing, or mean waveform shape regardless of their varying features in the spectral or temporal domain (though see: Duchet et al., 2021; Enz et al., 2021; Zich et al., 2020).

Here, we present a novel adaptive method for single-trial burst detection that captures the entire range of burst amplitudes, and apply it to high precision magnetoencephalography (MEG) sensor data. In line with previous findings, the overall rate of bursts detected with this method closely tracks changes in average beta amplitude, but bursts are widely diverse in time-frequency space. Using a biophysical model of burst generation, we show that variations in the timing, strength, and duration of the deep and superficial layer drives all change burst duration, peak amplitude, peak frequency, and frequency span via modulations of the cumulative dipole waveform shape generated by the model. Burst shape, and thus the underlying generating mechanism, is therefore not uniquely identified by features in the TF domain. We then show that beta bursts in human MEG data indeed occur in a wide range of waveform motifs, only some of which are predicted from the model, and which deviate greatly from the mean burst waveform. Finally, we show that bursts with different waveforms, unpredicted by the model, are differentially rate-modulated during a visuomotor task, and therefore likely have different functions. These results thus serve as a novel demonstration that beta bursts in human sensorimotor cortex are not unitary, and that treating them as a homogeneous signal risks overlooking a potentially rich source of information about their mechanisms and functional roles.

## Method

### Behavioral task

Thirty-eight healthy, right-handed, volunteers with normal or corrected-to-normal vision and no history of neurological or psychiatric disorders participated in the experiment (25 female, aged 20 - 35 years, M = 26.69, SD = 4.11 years). The study protocol was in accordance with the Declaration of Helsinki, and all participants gave written informed consent which was approved by the regional ethics committee for human research (CPP Est IV - 2019-A01604-53).

Participants completed a cued visuomotor adaptation task in which they made rapid, joystick-based reaching movements to a visually presented target. At the start of each trial, subjects were required to visually fixate on a small (0.6° × 0.6°) central target which was a combination of a bullseye and crosshairs (Thaler et al., 2013). The joystick controlled the position of a small (0.5 × 0.5°) square white cursor. Following a variable delay (1 - 2 s), a circular random dot kinematogram (RDK) was presented, with coherent motion in either the clockwise or counter-clockwise direction. On the outer edge of the RDK, five circular potential reach targets appeared in 30° increments from −120° to 0°, relative to the fixation point. On each trial, one of these targets (either at −90°, −60°, or −30°) was larger (3.25°) and green, indicating that the participant would have to reach for that target after the go cue, and the others were smaller (1.625°) and gray colored. The RDK disappeared after 2 s, after which only the gray potential reach targets and the green instructed target were visible. After a variable delay period (0.5 - 1 s), the gray targets disappeared, leaving only the green instructed target. This served as the go cue, instructing the participant to use the joystick to rapidly reach for this target. Trials ended once the distance between the cursor and the fixation target exceeded that of the center of the instructed target (7°), if the reach was started too early (the distance to the fixation point exceeded 1° before the disappearance of the gray potential targets), or if the reach was not completed quickly enough (1 s after the go cue including reaction time and reach duration, with a reach duration of less than 500 ms). Each trial was separated by an inter-trial interval of random duration (1.5 - 2 s), during which participants were required to bring the cursor back to the fixation target. Once the inter-trial interval duration had passed and the cursor was within 1° of the fixation target, the next trial began.

Participants were split into 2 groups. On each trial, the RDK consisted of various levels of clockwise or counter-clockwise coherent motion, or no coherent motion at all, and the visual location of the cursor was rotated by −30°, 0°, or 30°, depending on the participant group and trial condition. For the explicit group (N = 20), the visuomotor rotation followed the direction of coherent motion of the RDK (−30° for counterclockwise coherent motion, 30° for clockwise, and no rotation for no coherent motion). Participants therefore could not adapt to the variable rotation, but could predict it from the RDK with varying levels of difficulty, and thus adjust their motor preparation during the delay period. The reaches of the implicit group were rotated by −30° in each trial, meaning there was no relationship between the direction of motion coherence in the RDK and that of the visuomotor rotation, but that participants could implicitly adapt to the constant rotation.

The RDK was presented within a 7° circular window centered on the fixation point with 200, 0.3° diameter white dots, each moving at 10°/s. On each trial, a certain percentage of the dots (specified by the motion coherence level) moved coherently around the center of the aperture in one direction, clockwise or counter-clockwise at a distance between 2° and 7° from the fixation target. The remaining dots moved in random directions, with a consistent path per dot. Dot lifetime was normally distributed (M = 416.67, SD = 83.33 ms). The levels were individually set for each participant by using an adaptive staircase procedure (QUEST; Watson & Pelli, 1983) to determine the motion coherence at which they achieved 82% accuracy in a block of 40 trials at the beginning of each session. During this block, participants had to simply respond with the left or right button to counterclockwise or clockwise motion coherence. The resulting level of coherence was then used as low, and 150% and 200% of it as medium and high, respectively.

Participants first completed 1-3 short training blocks of 12 trials, during which the RDK had no coherent motion in any direction, and there was no visuomotor rotation. During these blocks, the location of the joystick-controlled cursor was visible for the entire duration of the trial (online feedback), the cursor turned red once the reach distance exceeded the extent of the center of instructed target (7°), and a short message was shown for 2 s telling subjects if they began the reach too early, did not complete the reach fast enough, or were too far (> 1.625°) from the center of the target. Once participants correctly completed at least 75% of the trials within a training block, they performed one block of 56 trials which also contained no coherent RDK motion and no visuomotor rotation, but the cursor disappeared once its distance from the fixation target exceeded 1° and reappeared and turned red once it reached the extent of the center of the instructed target (endpoint feedback). At the end of each trial in this block, a short message was also shown for 2 s giving feedback on the timing and accuracy of the reach. Each group of participants then completed 7 blocks of 56 trials each with visuomotor rotation. In each of these trials, the RDK had zero, low, medium, or high levels of coherent motion in the clockwise or counterclockwise direction. For the explicit group, the visuomotor rotation was 0, −30, or 30°, depending on the direction of coherent motion in the RDK. For the implicit group, the visuomotor rotation was always 30°, regardless of the direction of coherent motion in the RDK. Finally, both groups completed one washout block of 56 trials in which there was no coherent motion and no visuomotor rotation. For the purposes of this study, data were aggregated across participant groups, blocks (except for the training blocks), and trial conditions. The task was implemented in Psychtoolbox (v3.0.16; Brainard, 1997; Pelli, 1997) and run using Matlab R2017b.

### MRI acquisition and head-cast construction

Prior to MEG data acquisition, each participant underwent an MRI session using a 3T Siemens Sonata system (Erlangen, Germany). A T1-weighted scan was acquired using a magnetization-prepared rapid gradient-echo (MPRAGE) pulse sequence with 1 mm isotropic voxel size (256 × 256 × 256 voxels), a repetition time (TR) of 2100 ms, an echo time (TE) of 3.33 ms, inversion time (TI) of 900 ms, and a grappa factor of 3. For the co-registration of the MRI and the MEG data, vitamin E tablets were placed at the nasion and the left and right ear canals.

We used the 1 mm T1 MRI volumes of the subjects in order to construct an individualized foam head-cast for each participant in order to reduce between-session co-registration error and within-session head movement (Bonaiuto et al., 2018; Meyer et al., 2017). The scalp surface was extracted from the T1 volume using Freesurfer (v6.0.0; Fischl et al., 2002) and used as a mold for the inner surface of the head-cast, with the outer surface defined by a 3D model of the MEG dewar. The surface models were then positioned relative to a 3D model of the MEG dewar using Rhinoceros 3D (https://www.rhino3d.com) in order to minimize the distance between the scalp and the sensors without obstructing the participant’s view. The resulting model was then printed using a Raise 3D N2 Plus 3D printer (https://www.raise3d.com). The 3D printed model was placed inside a replica of the MEG dewar, and the space between the head model and the dewar replica was filled with polyurethane foam (Flex Foam-it! 25; https://www.smooth-on.com) to create the participant-specific head-cast into which the fiducial coils were placed during scanning.

### MEG acquisition and preprocessing

MEG data were acquired using a 275-channel Canadian Thin Films (CTF) MEG system with superconducting quantum interference device (SQUID)-based axial gradiometers (CTF MEG Neuro Innovations, Inc. Coquitlam, Canada) in a magnetically shielded room. Participants were in the supine position during the recording. A video projector (Propixx VPixx, VPixx Technologies Inc., Canada) was used to display visual stimuli on a screen with a refresh rate of 120Hz (∼80 cm from the participant), and a joystick (NATA Technologies, Canada) was used for participant responses. The data collected were digitized continuously at a sampling rate of 1200 Hz. Eye movement data was collected using Eyelink 2000 eye tracker (SR Research, Ontario, Canada) which tracked monocular eye movements at 1000 Hz and was calibrated before the start of recording.

Data preprocessing was performed using the MNE-Python toolbox (Gramfort et al., 2014) unless stated otherwise. MEG data was downsampled to 600 Hz and filtered (low pass 120 Hz zero-phase FIR filter with a Hamming window). Line noise (50 Hz) was removed using an iterative version of the Zapline algorithm (de Cheveigné, 2020) implemented in the MEEGKit package (https://nbara.github.io/python-meegkit/). using a window size of 20 Hz for polynomial fitting and 5 Hz for noise peak removal and interpolation. Ocular movement and cardiac related artifacts were isolated by running Independent Component Analysis (ICA, InfoMax, 25 components were extracted) on a copy of the MEG data (band pass filtered from 1 to 60 Hz), implemented in the scikit-learn library (Pedregosa et al., 2011). The eye tracking data was first cropped and resampled to match the MEG signal, and then components containing ocular artifacts were identified by correlating each of the 25 components with the horizontal and vertical eye movement signals. Blink detection was done by thresholding the vertical eye movement beyond the vertical resolution of the screen. Each component time course was then correlated (Pearson’s *r*) with the horizontal and vertical gaze position signals before and after removing blinks. Each component thus had 4 correlation coefficients, and was classified as an ocular movement artifact if all correlation coefficients were above *r* = 0.15, and the average correlation was above *r* = 0.25. Cardiac artifacts were identified by applying the ECG R peak detector (https://github.com/berndporr/py-ecg-detectors; Porr & Howell, 2019) to each component, and choosing the one with the lowest inter-peak temporal variance. The component choice was manually verified prior to removal from the original (prior to band pass filtering) dataset.

The data were then epoched around two events within each trial, between −1 and 2 s relative to the onset of the visual stimulus (visual epochs), and between −1 and 1.5 s relative to the end of the reaching movement (motor epochs). We analyzed data from 11 sensors above the left sensorimotor area, contralateral to the hand used to make the movement (Figure 2A: inlay).

### Burst detection

We developed a novel, adaptive burst detection algorithm to ensure that all potentially relevant burst events were detected across a wide range of beta amplitudes (Figure 1). The algorithm operates iteratively on single trial TF decompositions, and continues until no more bursts are detected in the trial. The estimated aperiodic spectrum (Figure 1A) is first subtracted from each single trial TF decomposition (Figure 1B) and iterative burst detection then operates on the residual amplitude (Brady & Bardouille, 2022). On each iteration, the algorithm detects the global maximum amplitude in TF space, and fits a two-dimensional Gaussian to this peak by computing the symmetric full-width at half maximum (FWHM) in the time and frequency dimensions (Figure 1C). This 2D Gaussian parametrization thus defines burst features in TF space: peak time, duration, peak amplitude, peak frequency, and frequency span. The Gaussian is then subtracted from the TF decomposition, and the next iteration operates on the resulting residual TF matrix. This process continues until there are no global maxima above the noise floor remaining (2 standard deviations above the mean amplitude over all time and frequency bins, recomputed on each iteration). To avoid edge effects near the limits of the beta band, we applied the algorithm to TF data between 10 and 33 Hz, but only bursts with a peak frequency within the beta band (13 - 30 Hz) were retained for further analysis.

**Figure 1.**
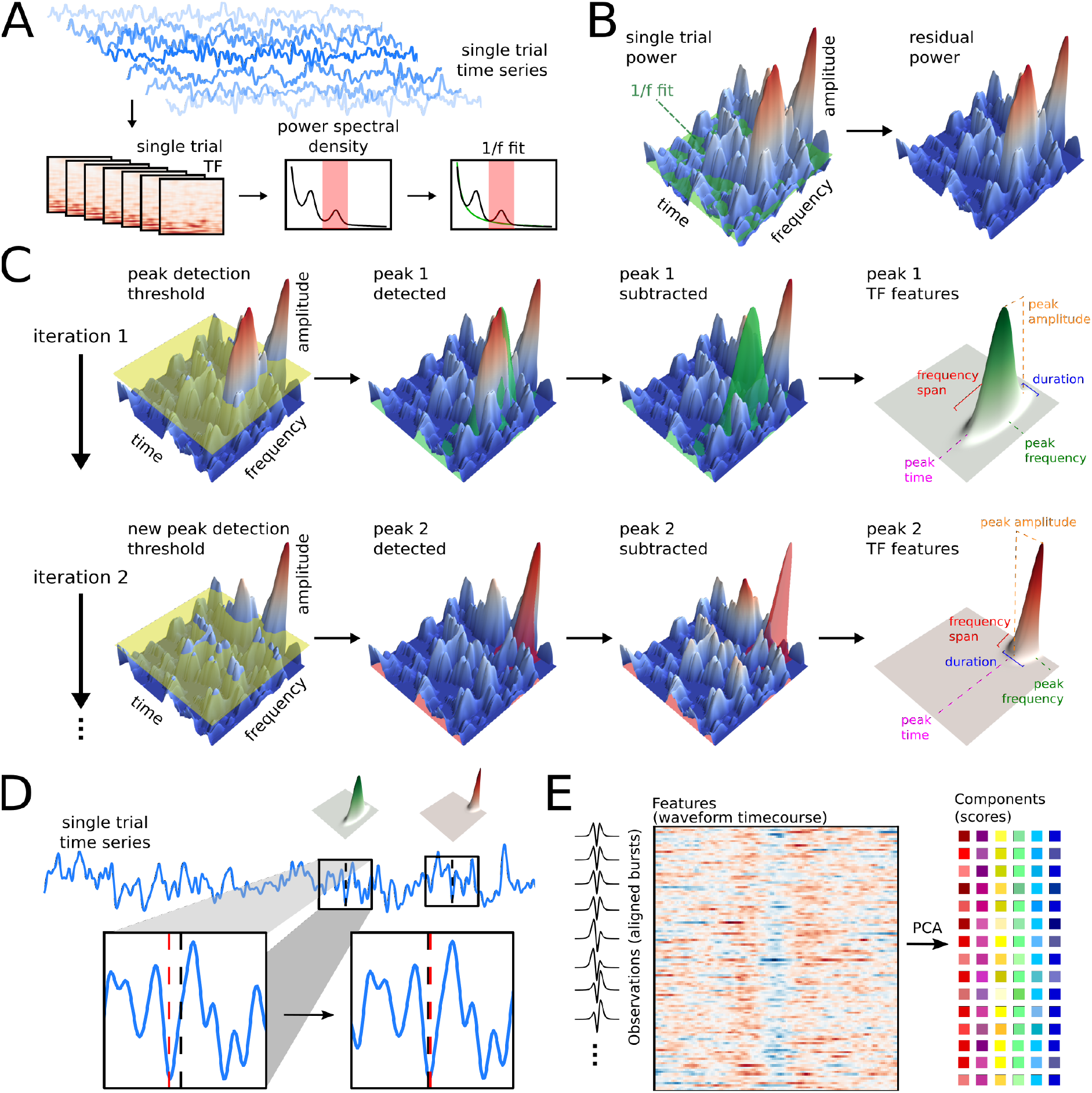
Iterative burst detection and waveform analysis. (A) The MEG single-trial signal time course was decomposed into the time-frequency (TF) domain using superlet transformation. The power spectral density (PSD) was calculated per trial by averaging the TF power over the time dimension, and then averaged over trials. A 1/f function was fitted to the averaged PSD to account for aperiodic neural activity. (B) The aperiodic fit was then subtracted from each single trial TF in order to isolate bursts with amplitude above that of background neural activity and noise. (C) After removing the aperiodic influence, bursts from each single trial TF were detected using an iterative algorithm. During each iteration, the global maximum was identified, defining the peak time and frequency, and a two-dimensional Gaussian was fitted to the peak to compute burst duration and frequency span. The Gaussian was then subtracted from the single trial TF, and the following iteration operated on the residual. The process terminated when there were no peaks above the noise floor (here set to 2 standard deviations of the single trial TF on each iteration). (D) For each burst identified from the single trial TFs, a segment of 260 ms was extracted from the corresponding trial time series, centered on the peak time. The peak time was then adjusted to the closest time point with zero phase after band pass filtering within the burst’s frequency span. (E) PCA was then applied to the phase-aligned (but not band pass filtered) burst waveforms, resulting in a score for each burst along each of 20 principal components.

We used the superlet transform (Moca et al., 2021) to compute single-trial TF decompositions. The superlet transform is a relatively new method of TF decomposition that more optimally balances time and frequency resolution than other commonly used methods like the short-time Fourier transform or continuous wavelet transform, making it especially well suited to detect transient bursts in restricted frequency bands. We used an adaptive superlet transform based on Morlet wavelets with varying central frequency (1 - 120 Hz) and number of cycles (4 cycles) under a Gaussian envelope. The order, which is a multiplier of the amount of cycles in the wavelet, was linearly varied from 1 to 40 over the frequency range. The TF decomposition was then used to calculate the power spectral density (PSD) for each sensor by averaging single-trial TF power over time to obtain single trial PSDs, and then averaging PSDs over trials within each experimental block (56 trials; Figure 1A). We then used specparam (Donoghue et al., 2020) to parameterize the PSD of each sensor for each block and thus estimate the aperiodic spectrum using a linear function (log-log space equivalent of an exponential function, without a ‘knee’).

Based on their peak time, we then extracted the waveform for each detected burst from the “raw” time series (unfiltered, except for the 120 Hz low pass filter applied during preprocessing). To remove the effect of slower event-related field (ERF) dynamics on burst waveforms, the epochs were first averaged in the temporal domain to compute the ERF, and this was then regressed out of the signal for each trial. To determine the width of the time window for waveform extraction, we computed lagged coherence over all trials and sensors from 5 to 40 Hz and 2.5 to 5 cycles (Fransen et al., 2015). We used overlapping time windows with lag- and frequency-dependent widths, and Fourier coefficients were obtained for each time-window using a Hann windowed Fourier transform. The time window for waveform extraction was then defined as the FWHM of lagged coherence averaged within the beta band, yielding 5.25 cycles, or 255.81 ms at the mean beta frequency of 21.5 Hz (Figure S1), which we rounded up to 260 ms. The time series in this 260 ms window centered on the peak time was therefore extracted from the trial time series. In order to find the signal deflection corresponding to the peak in amplitude, we aligned burst waveforms by band pass filtering them within their detected frequency span (zero-phase FIR with a Hamming window), computing their instantaneous phase using the Hilbert transform, and recentering the “raw” (prior to band pass filtering) waveform around the phase minimum closest to the peak time detected in TF space (Boto et al., 2022; Figure 1D). If this time point was greater than 30 ms away from the TF-detected peak time, the burst was discarded. The DC offset was then subtracted from the resulting waveform. Finally, due to uncertainty in the orientation and source location of the dipoles generating measured sensor signals, we reversed the sign of burst waveforms in which the central deflection was positive (S. R. Jones et al., 2009). Open source code for the burst detection algorithm is available at https://github.com/danclab/burst_detection.

All measures of burst rate and mean beta amplitude over time were baseline corrected as the percent change from the mean value from 500 to 250 ms before the onset of the visual stimulus. The number of bursts detected per trial was compared between the visual and motor epochs using R (v4.2.0, R REF) with a generalized linear model with a Poisson distribution and log link function, with epoch type as a fixed effect and subject-specific offsets as random effects (lme4 v1.1.29; REF). The effect of epoch type was then assessed using a type II Wald *X*^2^ test (car v3.1.0; REF). Burst features in TF space (duration, peak amplitude, peak frequency, and frequency span) were compared between the visual and motor epochs using two-sample Kolmogorov - Smirnov tests (Pratt & Gibbons, 1981).

### Burst analysis

To classify the diversity of burst waveform shapes, principal component analysis (PCA, 20 components, implemented in the scikit-learn library (Pedregosa et al., 2011)) was applied to the phase aligned waveforms, with each waveform time point as a feature (Figure 1E). The principal components (PCs) were computed from a subset of the data, consisting of 20% of the waveforms, evenly sampled from each subject, block, and epoch, after removing waveforms outside of 10th - 90th percentile of median amplitude (*N* = 2,339,211). All detected bursts (*N* = 14,328,947), across subjects, blocks, and epochs, were then projected onto each PC, thus each burst received a score for each component representing the shape of its waveform along that dimension.

To determine which components were meaningful and not simply driven by noisy fluctuations of the neural field potential, a permutation approach was used (Vieira, 2012). To remove the correlation between the features (waveform time points), the matrix containing the data subset was shuffled within each time point (column) independently. PCA was then applied to the shuffled matrix using the same parameters as with an unshuffled subset. The p value for each PC was then given by the probability of the proportion of variance explained being lower after shuffling than for the unshuffled data. The higher this probability, the higher the proportion of explained variance contributed by the noise introduced by shuffling, thus indicating that the original component was mostly driven by noise. For each component, 100 permutations were run, using an alpha threshold of *p* = 0.05.

We then selected PCs that define dimensions along which the mean burst waveform shape varies across the visual and motor epochs. For each PC, the mean burst waveform score was calculated at each time point of the visual and motor epoch, and the variance of the mean score over the time course of each epoch was used for PC selection. Components with difference in the temporal variance of the mean score between epochs above a threshold of 0.002 were chosen for further analysis.

The mean baseline-corrected rate of bursts across the distribution of scores for each selected PC was calculated per subject. We used a one-sample cluster permutation test to determine significant deviation from the baseline. The family-wise error rate (FWER) was controlled by using a non-parametric resampling test with a maximum statistic (taken across all data points). As a statistic, the t-test with a variance regularization (“hat” adjustment (Ridgway et al., 2012), threshold p = 0.001) was chosen to minimize the effect of low variance data points, thus limiting spurious results. Threshold Free Cluster Enhancement (TFCE, starting threshold = 0, step = 0.2) was used to improve the statistical power of cluster detection, by employing an adaptive threshold on the level of a single data point (Smith & Nichols, 2009).

The distributions of burst duration, peak amplitude, peak frequency, and frequency span, were compared between bursts with scores from each quartile of the selected PCs using Bonferroni-corrected two-sample Kolmogorov - Smirnov tests (Pratt & Gibbons, 1981).

### Biophysical model

We used the open-source Human Neocortical Neurosolver (HNN) software to simulate a biophysical model of a cortical microcircuit driven by layer-specific synaptic inputs (HNN-core v0.2; https://hnn.brown.edu; Neymotin et al., 2020). This model has been fully described in prior publications (Law et al., 2022; Sherman et al., 2016; Shin et al., 2017). Here we used a slightly modified version of it, which was previously used to simulate beta bursts in the motor cortex (Bonaiuto et al., 2021). HNN’s underlying canonical neural circuit model simulates the generation of electrical currents in layered cortical columns that give rise to measurable EEG/MEG signals. These electrical currents (i.e., current dipoles), are assumed to be generated by post-synaptic, intracellular current flow in the spatially aligned dendrites of a large population of neocortical pyramidal neurons, and are simulated by HNN via the net intracellular electrical current flow in the pyramidal neuron dendrites, multiplied by their length (in nano-Ampere-meters). The net current dipole output is then scaled to fit the amplitude of recorded MEG data. The model contains multicompartment pyramidal neurons (PN) and single compartment interneurons (IN), located in infra- and supra-granular layers. Neurons receive excitatory synaptic input from simulated trains of action potentials in predefined temporal profiles that target the proximal apical/basal and distal apical dendrites of the PNs.

In the simulations described in this paper, we used a modified version of the default parameter set distributed with HNN that simulates beta bursts. In brief, this simulation contained 100 PNs and 35 INs per layer and received a proximal excitatory synaptic drive, simultaneous with a distal excitatory synaptic drive. The histogram of spikes from the inputs that generated these drives had a Gaussian profile. It has previously been shown that this pattern of inputs can generate bursts with the commonly observed beta burst waveform shape as part of the more continuous somatosensory mu-rhythm (S. R. Jones et al., 2009; Sherman et al., 2016), but here, we use the model to generate individual bursts only. Individual beta bursts occur when broad upward current flow from proximal inputs is synchronously disrupted by downward stronger and faster current flow from distal inputs. Results varied across simulations with identical parameters due to the stochastic nature of the exogenous proximal and distal drives: the timing of the synaptic drives were chosen from a Gaussian distribution. In the base simulations, the distribution of proximal drive inputs had a standard deviation of 20 ms, whilst that of the distal drive inputs had a standard deviation of 7 ms. For the proximal drive, the synaptic weight to supra- and infra-granular PNs was 3.5e-5 μS, lower than that of the distal drive (6e-5 μS). The mean delay between the proximal and distal drive was 0 ms. We ran five sets of simulations in order to ascertain how varying the timing, strength, and duration of the proximal and distal drives determine the beta waveform shape: 1) relative timing of the two drives (from −25 ms to 25 ms), 2) distal drive synaptic weight (from 4.5e-5 μS to 8.5e-5 μS), 3) proximal drive synaptic weight (from 0.5e-5 μS to 5.5e-5 μS), 4) distal drive duration (from 1 ms to 15 ms), and 5) proximal drive duration (from 10 ms to 40 ms). These parameter ranges were chosen in order to ensure that the resulting burst peak frequency was within the beta range, and that the bursts were driven by subthreshold dynamics (S. R. Jones et al., 2009; Sherman et al., 2016). Each simulation type was run with 10 different parameter values, equally spaced within the range tested, and for 500 trials per value, with each trial (200 ms) resulting in a single burst. The superlet transform (adaptive, 4 cycles, 100 equally spaced central frequencies between 2 and 40 Hz) was applied to the burst waveform from each trial, and the duration, peak amplitude, peak frequency, and frequency span were computed in TF space as per the burst detection algorithm described above. Relationships between model parameters and burst duration, peak amplitude, peak frequency, and frequency span were evaluated using growth curve models implemented in R (v4.2.0; R Core Team, 2022) using the lme4 library (v1.1.29; Bates et al., 2014). The parameter values were modeled with a second-order (quadratic) orthogonal polynomial, and polynomial significance was estimated using type II Wald F tests (car v3.1.0; Fox et al., 2019). Burst waveforms generated by the model were then scaled to match the bursts detected from the human MEG data (scaling factor = 9e^-16^) and projected onto each dimension defined by the PCA fit to the human subject data.

All preprocessing, analysis, and simulation code is available at https://github.com/maciekszul/DANC_beta_burst_PC_analysis.

## Results

### Beta bursts are diverse

We analyzed data recorded from a cluster of left central MEG sensors while subjects performed a cued visuomotor task involving presentation of a visual stimulus indicating the target location, a variable delay, followed by a right-handed reaching movement made with a joystick to the target. We detected beta bursts in the 13 - 30 Hz range using a novel burst detection algorithm (see Methods), and extracted burst waveforms from two trial epochs: one aligned to the onset of the visual stimulus (visual epochs), and one aligned to the offset of the reaching movement (motor epochs). In line with previous work (Little et al., 2019), the overall burst rate decreased following the onset of the visual cue, further decreased during the reaching movement, and increased again following the end of the reach, closely matching changes in mean beta power (Figure 2A). More bursts were therefore detected during the visual compared to the motor epoch (visual: *M* = 477.45, *SD* = 83.07 bursts per subject per trial; motor *M* = 392.83, *SD* = 68.92 bursts per subject per trial; *X*^2^(1) = 135,479, *p* < 0.001). Across both epoch types, bursts were variable in terms of time-frequency features, but there were no differences between epochs in distributions of burst duration (*D* = 0.008, *p* < 0.001), peak amplitude (*D* = 0.01, *p* < 0.001), peak frequency (*D* = 0.004, *p* < 0.001), or frequency span (*D* = 0.006, *p* < 0.001). In general, bursts were short (overall *M* = 185 ms, *SD* = 97 ms; Figure 2B), burst peak amplitude distributions had a long tail (overall *M* = 18.91, *SD* = 13.14 fT; Figure 2C), burst peak frequency was centered around 21 Hz (overall *M* = 21.35, *SD* = 4.76 Hz; Figure 2D), and bursts spanned 1 to 4 Hz in the frequency domain (overall *M* = 2.01, *SD* = 0.81 Hz; Figure 2E). As described previously (Bonaiuto et al., 2021; Kosciessa et al., 2020; Little et al., 2019; Sherman et al., 2016), the median burst waveform shape was wavelet-like (Figure 2F), with a prominent central negative deflection, symmetrically surrounded by positive deflections on either side. Individual burst waveforms, however, deviated greatly from the median with a relatively low signal-to-noise ratio (SNR) outside of this central negative deflection and surrounding peaks (overall *M* = −27.16, *SD* = 14.60 dB; Figure 2F).

**Figure 2.**
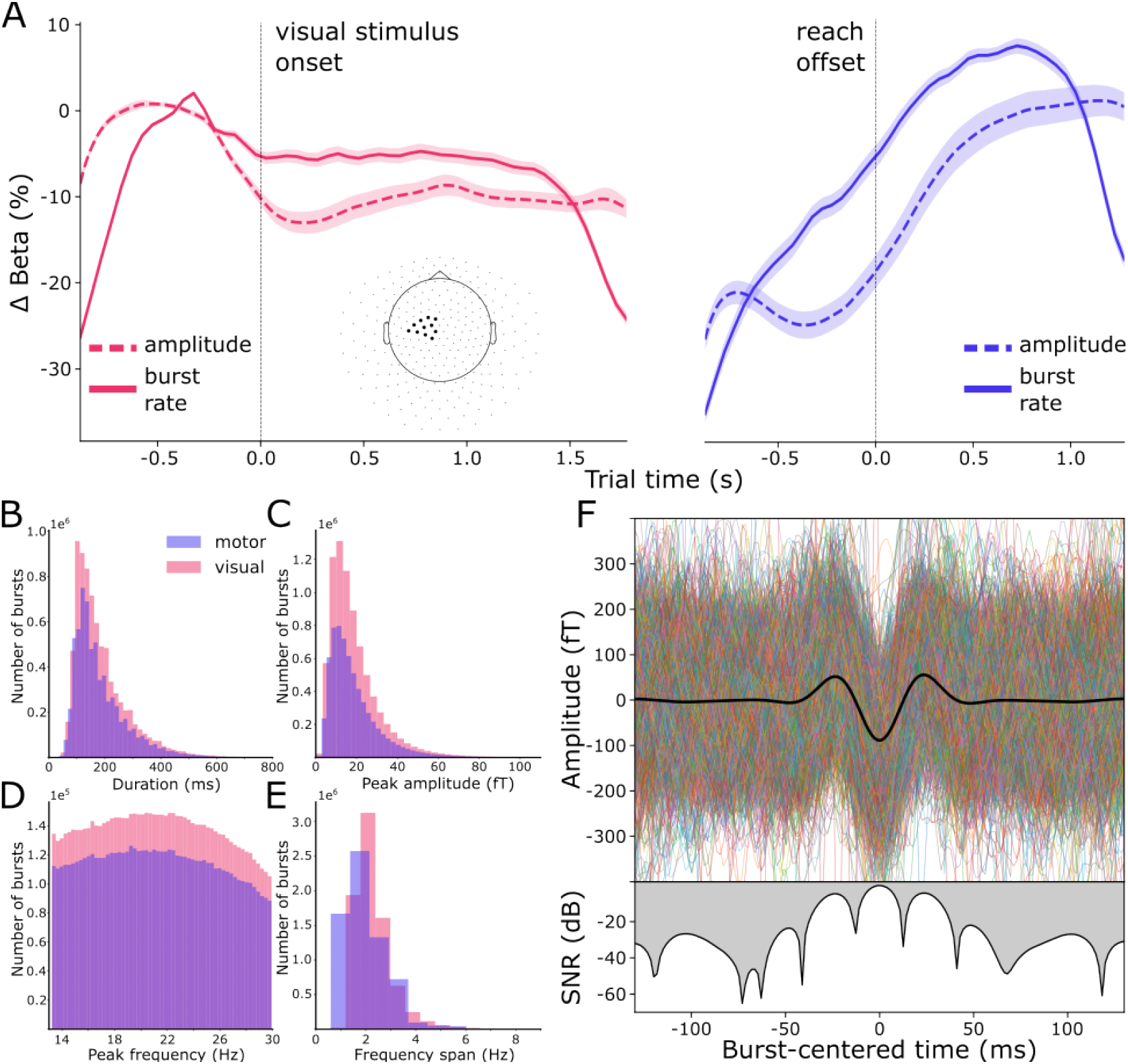
Burst dynamics and summary of burst features. (A) The median burst rate (solid line; shaded area shows the SEM) decreased following the onset of the visual stimulus, further dropped during the movement, and rebounded following the end of the movement. Burst rate dynamics closely matched those of median beta amplitude (dashed line; shaded area shows the SEM). The inset shows the 11 sensors used in this analysis, located above sensorimotor areas contralateral to the hand used. (B-E) The distributions of burst peak duration (B), peak amplitude (C), peak frequency (D), and frequency span (E) were similar in the two phases of the task. (F) The median aligned waveform (thick black line, top) over all detected bursts had a wavelet-like shape, but there was great variability in the waveforms of individual bursts (thin colored lines, top). The SNR of burst waveforms was highest around the central negative deflection and surrounding peaks, but less than −20 dB for the first and last 100 ms.

### Burst variability can be explained by biophysical model parameters

One of the most prominent biophysical models of beta burst generation simulates bursts as being driven by temporally aligned proximal and distal drives to the deep and superficial cortical layers (Figure 3A; Sherman et al., 2016). The model predicts that the beta burst waveform shape is caused by the aggregate of these two, oppositely oriented, current flows. The proximal drive targets the deep layers and consists of a temporally dispersed, weak excitatory synaptic input, whilst the distal drive targets the superficial layers and is stronger, but briefer. The temporal alignment of these drives results in cumulative dipole moments with wavelet-like waveform shapes which match the median burst waveform shape observed in the MEG data, but not the variance around the median (Figure 3B). TF decomposition of the cumulative dipole moment generated by the model reveals a transient increase in beta amplitude, from which the duration, peak amplitude, peak frequency, and frequency span can be computed (Figure 3C). However, the individual burst waveforms generated by the base model have a wide range of durations (*M* = 130.11, *SD* = 4.10 ms), peak amplitudes (*M* = 61.68, *SD* = 17.69 nAm), peak frequencies (*M* = 15.52, *SD* = 2.34 Hz), and frequency spans (*M* = 8.46, *SD* = 1.30 Hz; Figure 3D) in the TF domain. The model therefore generates variable burst waveforms which result in variability in TF-based features of individual bursts, even with a single set of distal and proximal synaptic drive parameter values.

**Figure 3.**
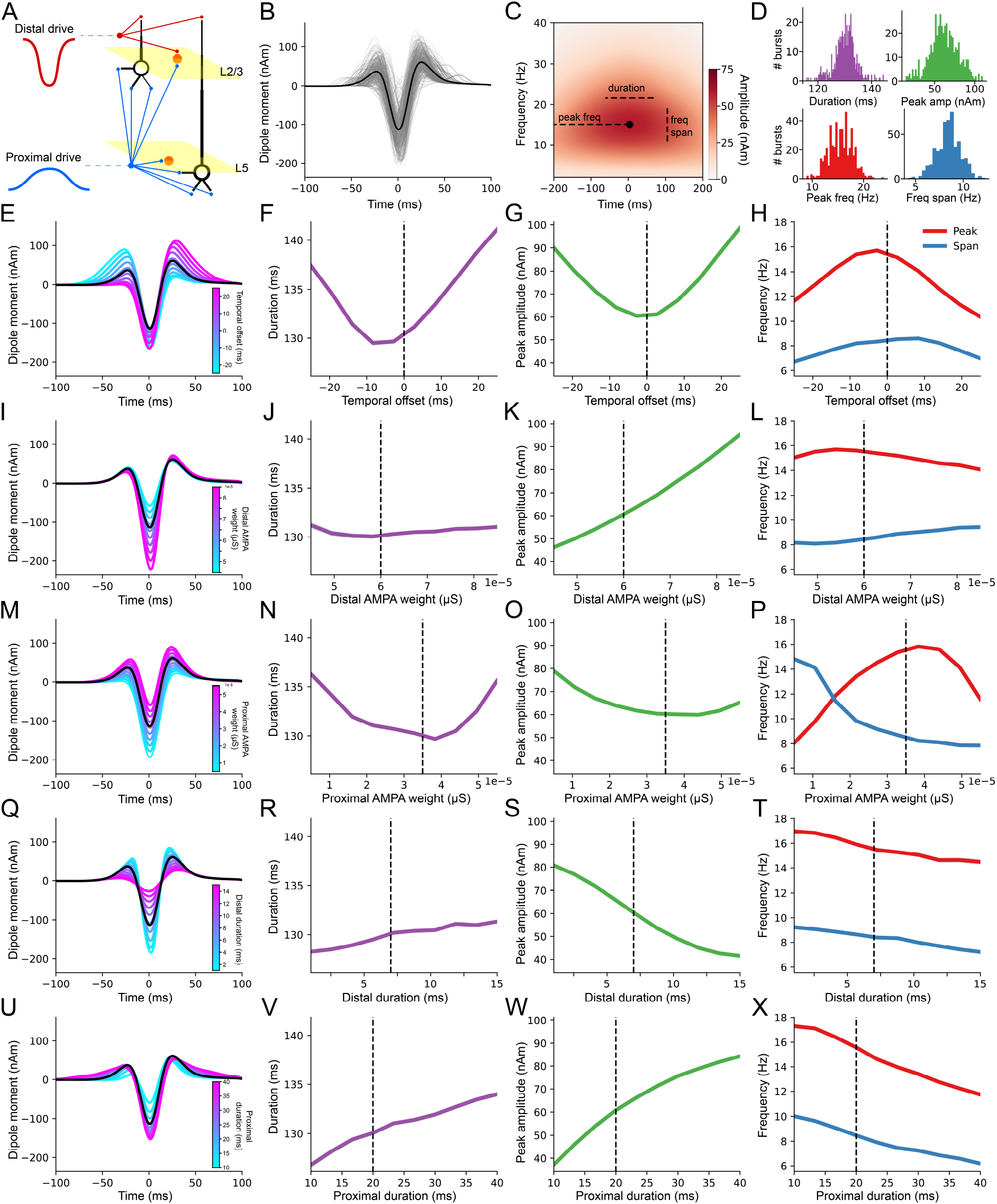
The biophysical model can generate bursts with different waveform shapes. (A) The biophysical model of beta burst generation consists of a proximal (infragranular) drive, and a distal (supragranular) drive. The Gaussians represent the duration of each drive and their polarity represents the direction of intracellular current flow. The model consists of multiple-compartment pyramidal neurons (black), and local inhibitory interneurons (orange). (B) The combination of proximal and distal drives generates a cumulative dipole moment that closely matches the median burst waveform observed in human MEG data (gray lines are individual burst waveforms generated from the base model, black line is the average waveform). (C) The mean TF spectrum over all generated bursts from the base model. The time and frequency at which the peak amplitude (black dot) is detected are used to compute the burst frequency span and duration (dashed lines). (D) The model generates bursts with a range of durations, peak amplitudes, peak frequencies, and frequency spans. (E) When the peak of the proximal drive is offset between −25 and 25 ms, the model generates burst waveform shapes with varying degrees of asymmetry. (F-H) Waveform asymmetry has nonlinear effects on mean burst duration (F), peak amplitude (G), peak frequency (H), and frequency span (H). The vertical dashed lines indicate the proximal drive offset in the base model (0 ms). (I) The magnitude of the central negative deflection in the burst waveform is modulated by the strength of the distal drive. (J-L) As in F-H, for the strength of the distal drive. (M) The strength of the proximal drive jointly shifts the amplitudes of the central negative and surrounding positive deflections of the waveform. (N-P) As in F-H, for the strength of the proximal drive. (Q) The duration of the distal drive modulates the amplitude and sharpness of the central negative and surrounding positive deflections in the waveform shape. (R-T) As in F-H, for the standard deviation of the distribution of the distal drive timing. (U) The duration of the proximal drive modulates the duration of the surrounding positive deflections and the magnitude of the central negative deflection in the burst waveform. (V-X) As in F-H, for the standard deviation of the distribution of the proximal drive timing.

In order to determine if TF-based burst features could, in principle, distinguish between burst waveform shapes, we simulated bursts across a range of model parameter values. The most influential model parameters in generating the stereotypical waveform shape are the relative timing, strength, and duration of the two synaptic drives. We therefore ran the model using a range of proximal input timing (changing the relative timing of the two drives), distal and proximal input strength (AMPA synapse weights), and durations (standard deviations of the drive timing distribution). Varying the timing of the proximal input peak, from 25 ms before to 25 ms after the distal input peak, generated a spectrum of waveform shapes that varied in their asymmetry (Figure 3E). Waveform asymmetry had nonlinear effects on burst duration (linear *F*(1) = 609.84, *p*< 0.001, quadratic *F*(1) = 2194.27, *p*< 0.001; Figure 3F), peak amplitude (linear *F*(1) = 150.01, *p*< 0.001, quadratic *F*(1) = 2624.43, *p*< 0.001; Figure 3G), peak frequency (linear *F*(1) = 336.66, *p*< 0.001, quadratic *F*(1) = 2242.49, *p*< 0.001; Figure 3H), and frequency span (linear *F*(1) = 68.07, *p*< 0.001, quadratic *F*(1) = 1043.28, *p*< 0.001; Figure 3H). Running the model with a range of distal input strengths altered the magnitude of the central negative deflection (Figure 3I), which had a nonlinear effect on burst duration (quadratic *F*(1) = 16.15, *p*< 0.001; Figure 3J), peak amplitude (linear *F*(1) = 4216.58, *p*< 0.001, quadratic *F*(1) = 42.01, *p*< 0.001; Figure 3K), and peak frequency (linear *F*(1) = 105.04, *p*< 0.001, quadratic *F*(1) = 41.23, *p*< 0.001; Figure 3L), and a linear effect on frequency span (*F*(1) = 389.86, *p*< 0.001; Figure 3L). Varying the strength of the proximal synaptic drive jointly shifted the magnitude of the central negative and surrounding positive burst waveform deflections (Figure 3M). This had a nonlinear effect on the burst duration (linear *F*(1) = 36.40, *p*< 0.001, quadratic *F*(1) = 617.82, *p*< 0.001; Figure 3N), peak amplitude (linear *F*(1) = 458.50, *p*< 0.001, quadratic *F*(1) = 408.73, *p*< 0.001; Figure 3O), peak frequency (linear *F*(1) = 867.32, *p*< 0.001, quadratic *F*(1) = 1299.89, *p*< 0.001; Figure 3P), and frequency span (linear *F*(1) = 4746.92, *p*< 0.001, quadratic *F*(1) = 865.55, *p*< 0.001; Figure 3P). Differences in the temporal dispersion of the two synaptic drives had more complex effects on burst waveform shape. The duration of the distal drive changed the amplitude and sharpness of the central negative and surrounding positive waveform deflections (Figure 3Q), which had a linear effect on burst duration (*F*(1) = 149.79, *p*< 0.001; Figure 3R), and nonlinear effects on burst peak amplitude (linear *F*(1) = 3747.90, *p*< 0.001, quadratic *F*(1) = 53.94, *p*< 0.001; Figure 3S), peak frequency (linear *F*(1) = 310.70, *p*<0.001, quadratic *F*(1) = 7.11, *p* = 0.008; Figure 3T), and frequency span (linear *F*(1) = 876.60, *p*< 0.001, quadratic *F*(1) = 4.38, *p* = 0.037; Figure 3T). The duration of the proximal drive changed the duration of the surrounding positive deflections and the magnitude of the central negative waveform deflections (Figure 3U). This had nonlinear effects on duration (linear *F*(1) = 817.31, *p*< 0.001, quadratic *F*(1) = 13.12, *p*< 0.001; Figure 3V), peak amplitude (linear *F*(1) = 4112.50, *p*< 0.001, quadratic *F*(1) = 166.49, *p*< 0.001; Figure 3W), and frequency span (linear *F*(1) = 3026.30, *p*< 0.001, quadratic *F*(1) = 40.20, *p*< 0.001; Figure 3X), and a linear effect on peak frequency (*F*(1) = 2025.90, *p*< 0.001; Figure 3X). Each model parameter tested resulted in nonlinear changes in nearly all burst features defined in TF space, thus preventing inference of underlying neural circuit dynamics by inspection of TF burst features alone. In order to explore beta burst variability with respect to underlying mechanistic models of burst generation, it is therefore necessary to analyze burst variability in the temporal, rather than TF, domain.

### PCA-derived waveform motif spectrums reveal shape specific task modulation

Having shown that varying the parameters of the drives to the model can result in a range of generated waveform shapes and that these waveforms have complex relationships with TF-based burst features, we then focused our analysis on the waveforms of bursts extracted from the human MEG sensor data. We applied PCA to these waveforms in order to identify motifs that explain variance in burst waveform shape, finding that 20 components explained 82% of the variance. We then ran a permutation test, shuffling the waveforms, in order to determine which components significantly contribute to variability in the observed waveforms. This revealed 18 significant components (*p*< 0.001; Figure 4A). We then further analyzed four components based on the temporal variance in mean burst score in the motor compared to the visual epoch, thus selecting dimensions along which the mean burst shape varied systematically over the course of the trial, and differently during the two epoch types. Each of these components defined dimensions along which the waveform shape varied markedly from the median waveform (Figure 4B-E; see Figure S2 for all 18 significant components). In each of these dimensions, the amplitude of peaks surrounding the central negative deflection, and that of the central deflection itself varied, but the most striking feature of each of the four components is that they represent waveforms with additional peripheral peaks. The mean burst waveform score for each of these components decreased following the onset of the visual stimulus, further decreased during the movement for components 8, 9, and 10, and then increased following the movement. Therefore, not only does the overall burst rate decrease pre-movement and increase post-movement, but the mean burst waveform shape also systematically changes over the course of the task.

**Figure 4.**
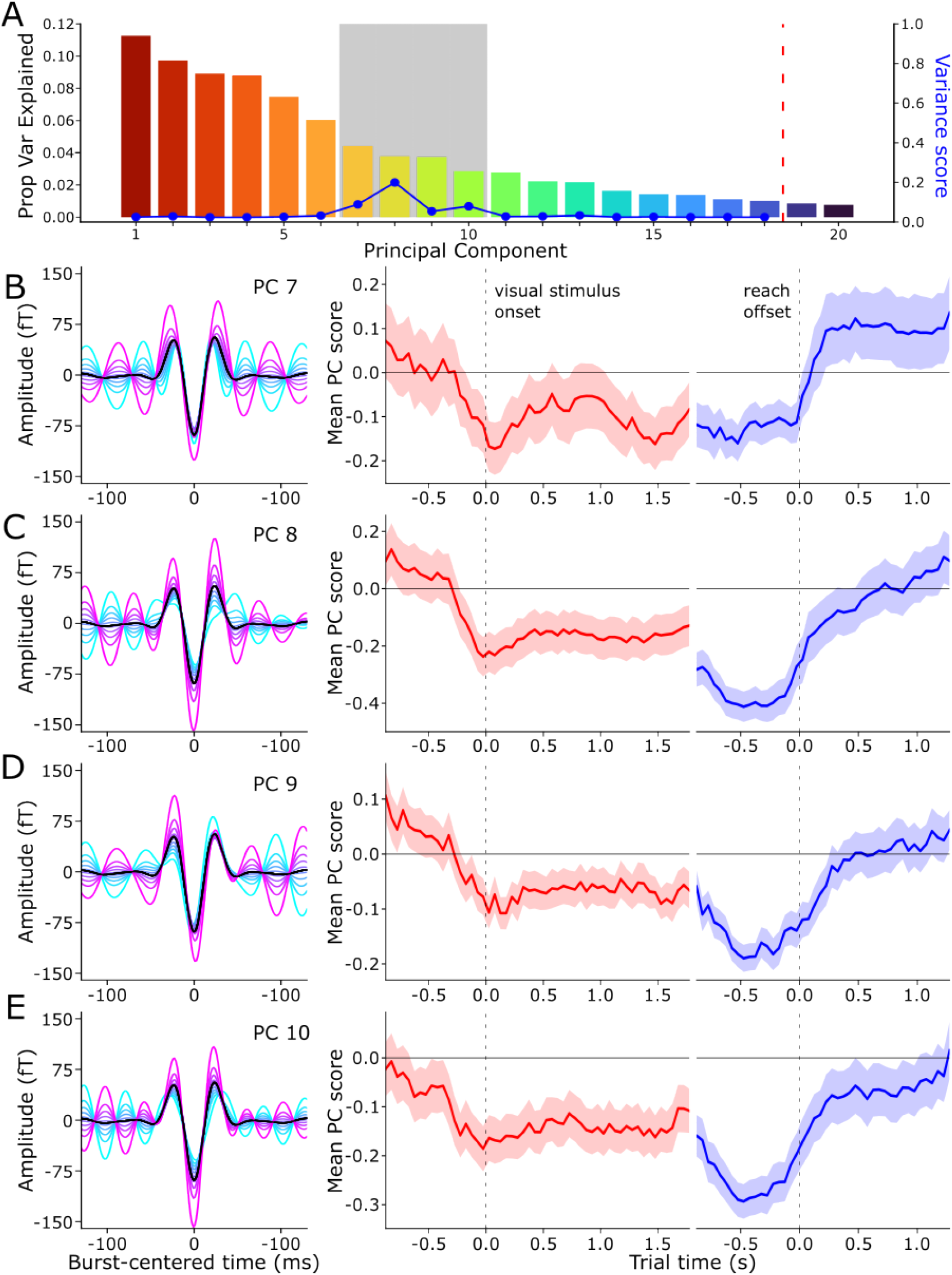
PCA reveals a variety of burst waveform motifs. (A) The histogram shows the proportion of the variance explained by each component. The vertical line shows the meaningful components used in further analyses as a result of the permutation analysis. The blue line overlaid on top of the histogram shows the variance score, which is the difference in the variance of a mean score time course between visual and motor epochs. The shaded rectangle highlights the components with a variance above threshold, which were then subsequently analyzed further. (B-E) For each chosen component, mean burst waveform shapes of bursts with a score within the 0 - 10th (cyan) to 90 - 100th (magenta) percentile of all burst scores for that component (left panels). The panels on the right show the time course of the mean burst score (solid line; shaded region shows SEM) for each component during the visual and motor epochs.

The movement-related changes in mean burst score for each of these components are confounded by burst rate. A reduction in mean burst score along any dimension could be due to a change in waveform shape with no change in burst rate, or some combination of a reduction in the rate of bursts with high scores, and an increase in the rate of those with low scores. For each selected component, we therefore examined how burst rate changed throughout the task, according to burst waveform shape along that dimension. With this aim, we binned bursts according to their component score, indicating their waveform shape, and the time during the trial in which they occurred. For each component score bin, we then baseline-corrected the burst rate using a time period prior to the onset of the visual cue, and then used a permutation test to determine significant deviations from the baseline (Figure 5; see Figure S3 for all significant components). Bursts with waveform shapes closest to the median waveform (i.e. those with scores close to 0 along each dimension) were the least modulated pre- and post-movement. Rather, for each selected component, bursts with higher scores (those in the fourth quartile) exhibited the greatest decrease in rate after the onset of the visual stimulus, followed by a transient increase, and then the greatest decrease in rate during the movement. The rate of bursts with scores in the second quartile along PCs 7, 9, and 10 did not change following the onset of the visual stimulus. Surprisingly, bursts with scores in the second percentile of PC 8 actually increased in rate after the visual stimulus onset and during the movement (Figure 5C-D), and those in the second percentile of PC 10 increased during movement (Figure 5G-H). The rate of bursts with scores in the first and third quartiles of each component only slightly decreased pre-movement and during movement. The post-movement period was marked primarily by an above-baseline increase in the rate of bursts with scores in the third and fourth quartile of PC 7 (Figure 5A-B), and the first and second quartiles of PCs 8, 9, and 10, and a return to baseline rate levels for bursts with scores in the fourth quartiles of PCs 8, 9, and 10. Bursts with different waveform shapes therefore exhibited diverse temporal dynamics, and differentially contributed to the classically observed pre-movement beta decrease and post-movement beta rebound signals.

**Figure 5.**
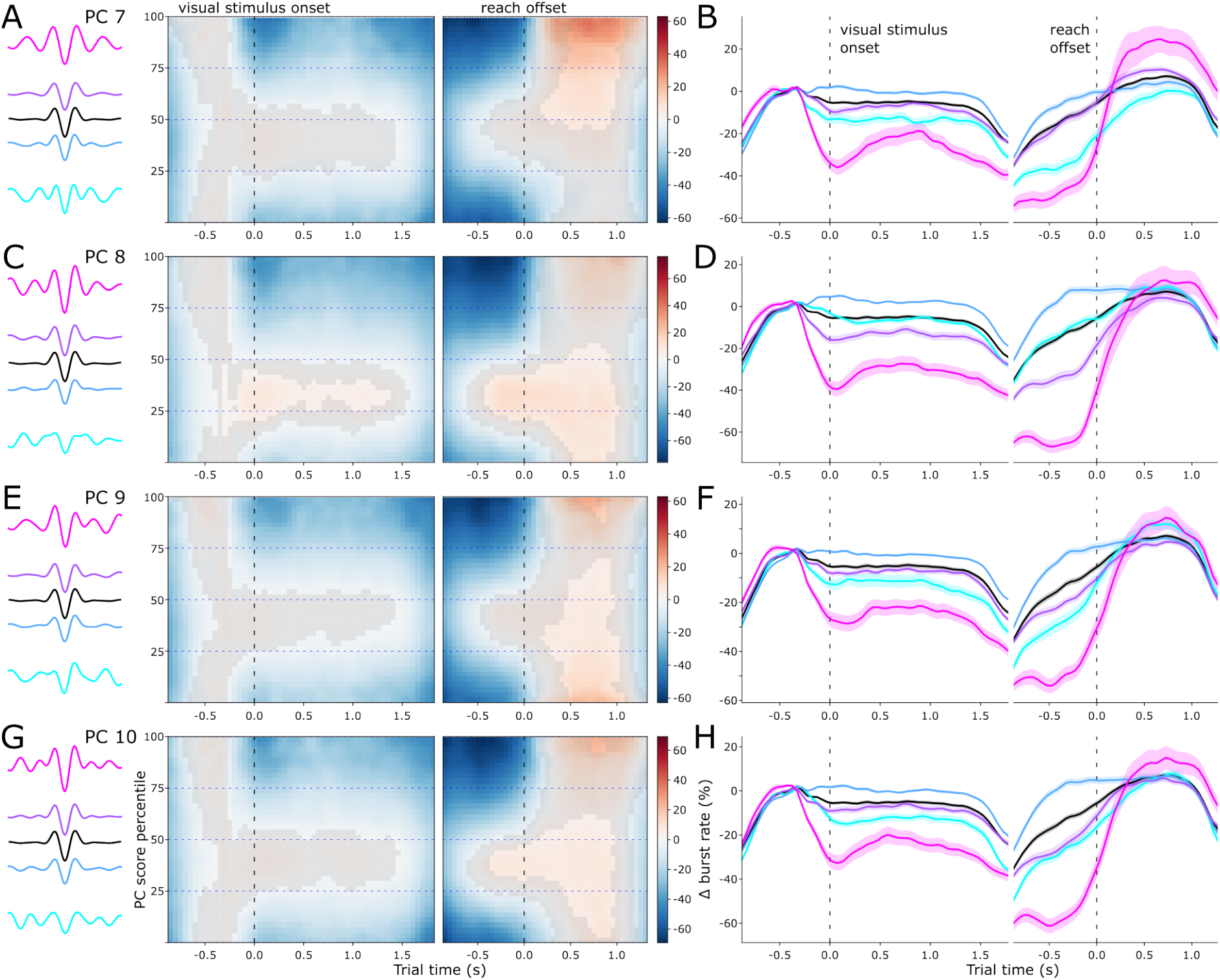
Burst motifs are differentially task-modulated. (A) The mean waveforms of bursts with scores in each quartile of the range of scores for PC 7 (left panel, colored lines) and the mean overall burst waveform (black), and the mean burst rate over time and across the range of PC 7 scores during the visual and motor epochs (right panel). The gray overlay obscures the time points and score ranges where the rate did not differ from the baseline rate. The horizontal dashed lines indicate the quartile limits. (B) The mean baseline-corrected rate of bursts with scores in each quartile of the range of scores along PC 7 (colored lines, the shaded area indicates the SEM) over the course of the visual and motor epochs. The black line represents the mean rate of all bursts (the shaded area indicates the SEM). (C-G) As in A-B, for PC 8 (C-D), PC 9 (E-F), and PC 10 (G-H).

Having demonstrated that specific waveform motifs differentially contribute to classic pre- and post-movement sensorimotor beta modulations, we then examined the distribution of TF-based features for each task-modulated motif. Burst scores for each PC were only weakly to moderately correlated with burst duration (PC 7: *ρ* = 0.12; PC 8: *ρ* = 0.10; PC 9: *ρ* = 0.04; PC 10: *ρ* = 0.10; all *p*< 0.001), peak amplitude (PC 7: *ρ* = 0.22; PC 8: *ρ* = 0.32; PC 9: *ρ* = 0.14; PC 10: *ρ* = 0.25; all *p*< 0.001), peak frequency (PC 7: *ρ* = −0.37; PC 8: *ρ* = 0.08; PC 9: *ρ* = 0.07; PC 10: *ρ* = 0.17; all *p*< 0.001), and frequency span (PC 7: *ρ* = 0.04; PC 8: *ρ* = 0.16; PC 9: *ρ* = 0.07; PC 10: *ρ* = 0.17; all *p*< 0.001; Figure S4). For each of the four selected PCs, the distributions of the TF-based features for each score quartile greatly overlapped (Figure S5). While bursts with scores in the 4th quartile each PC had higher mean amplitude than those in the first three quartiles (Table S1), the distributions of TF-based features were not different between quartiles for any of the task-modulated PCs (Table S2–5). Bursts with waveform shapes defined by these components could therefore not be distinguished by TF-based features alone. This underscores the utility of our approach, which adaptively detects all candidate burst events and then sorts them according to their underlying waveforms.

Finally, we sought to determine how well the biophysical model could capture the waveform variance described by each task-modulated waveform motif. We therefore projected the waveforms generated by the biophysical model onto the dimensions defined by the PCA fitted to the bursts detected in the human MEG data. For each set of simulations varying the temporal offset, distal and proximal drive strength, and distal and proximal duration, the parameter values were weakly to moderately correlated with nearly every principal component (Fig S6). No single parameter therefore modulated the burst waveform along one unique dimension. Of the four task-modulated burst waveform motifs, distal drive strength and duration were most correlated with PC 7 (strength: *ρ* = 0.43, *p*< 0.001; duration: *ρ* = −0.65, *p*< 0.001), PC 8 (strength: *ρ* = 0.54, *p*< 0.001; duration: *ρ* = −0.73, *p*< 0.001), and PC 10 (strength: *ρ* = 0.50, *p*< 0.001; duration: *ρ* = −0.76, *p*< 0.001), and temporal offset, distal strength, and distal drive duration were most correlated with PC 9 (temporal offset: *ρ* = −0.59, *p*< 0.001; distal strength: *ρ* = 0.44, *p*< 0.001; duration: *ρ* = −0.59, *p*< 0.001). The temporal offset between drives, and the strength and duration of the distal drive could, thus, explain some variance along these dimensions, but the range of mean model waveform component scores was very restricted compared to the human MEG data (temporal offset: 47.70 - 71.21 percentile of PC 9 scores; distal strength: 47.24 - 58.46 percentile of PC 7, 34.35 - 63.34 percentile of PC 8, 48.42 - 68.64 percentile of PC 9; 37.57 - 66.26 percentile of PC 10; distal drive duration: 35.79 - 56.72 percentile of PC 7, 20.22 - 62.27 percentile of PC 8, 40.94 - 67.06 percentile of PC 9, 21.03 - 68.07 of percentile PC 10; Figure S6). Whilst the model was able to explain some of the waveform variability described by the task-modulated burst motifs, it could not generate waveforms with shapes like those whose rate was most modulated pre- and post-movement.

## Discussion

We combined a novel burst detection algorithm, waveform analysis, and biophysical modeling to show that sensorimotor beta bursts occur with a wide range of waveform motifs which differentially drive movement-related changes in beta activity. In accordance with predictions from a biophysical model of somatosensory beta burst generation (S. R. Jones et al., 2009; Law et al., 2022; Neymotin et al., 2020; Sherman et al., 2016), we show that bursts have a wavelet-like mean shape, but individual bursts vary greatly from the mean. Variations in the timing, strength, and duration of the superficial and deep layer synaptic drives, according to the model, predict variability in bust waveforms that cannot be distinguished by TF-based features alone. We then applied dimensionality reduction to the waveform shapes of beta bursts detected over human sensorimotor cortex, and showed that bursts occur with a variety of waveform motifs. Finally, we have shown that the mean burst waveform changes systematically pre- and post-movement, and that this is caused by changes in the rate of bursts from specific motifs. Sensorimotor beta bursts are therefore not homogeneous events, and as a consequence, burst variability may provide the key to understanding how neural signals with TF-based signatures within the same frequency range can underlie the plethora of functional roles ascribed to “beta activity” (Engel & Fries, 2010; Kilavik et al., 2013; Little & Brown, 2014; Pfurtscheller et al., 1997; Reuter et al., 2022; Salenius & Hari, 2003).

The strategy employed by our burst detection method deviates from those commonly used in the field. A global threshold on beta amplitude or power is most commonly used, often based on a centrality measure (mean or median beta amplitude), calculated from the whole dataset (Bonaiuto et al., 2021; Brady & Bardouille, 2022; Diesburg et al., 2021; Enz et al., 2021; Feingold et al., 2015; Kehnemouyi et al., 2021; Little et al., 2019; Shin et al., 2017; Wessel, 2020; Zich et al., 2022), or a percentile of the beta power distribution (Anidi et al., 2018; Cagnan et al., 2019; Pauls et al., 2022; Sherman et al., 2016; Tinkhauser, Pogosyan, Tan, et al., 2017; Torrecillos et al., 2018; Yeh et al., 2020). The consequence of these approaches is that only high amplitude burst events are detected, neglecting low amplitude, but potentially informative bursts. For the TF data the most common methods are to use linear thresholds that are multiples of the mean or median beta amplitude, followed by various methods for local maxima detection (Brady & Bardouille, 2022; Diesburg et al., 2021; Enz et al., 2021; Shin et al., 2017; Wessel, 2020). More recently, based on efforts to parameterize aperiodic and periodic neural activity (Donoghue et al., 2020), a burst detection algorithm using a multiple of the aperiodic activity as a threshold has been introduced (Brady & Bardouille, 2022). Rather than using a fixed absolute threshold, our approach detects every peak above the aperiodic spectrum as a candidate burst event using an iterative algorithm to detect all bursts with amplitudes above the noise floor. It can therefore be seen as an extension of power spectra parameterization (Donoghue et al., 2020) from one dimensional PSDs to two dimensional TF decompositions. The result is that many more bursts across a wider range of amplitudes are detected, allowing subsequent analyses to determine which ones are functionally relevant.

Because of its general applicability and interpretability of results, we used PCA to characterize burst shapes. However, PCA creates a specific categorization of bursts by defining orthogonal components, thus yielding a Fourier-like decomposition of time series with components that appear as phase-shifted rhythmic deviations from the mean waveform at different frequencies. Moreover, because bursts can have negative scores along each dimension, the relationship between components and the underlying synaptic drives that generate burst waveforms is not easily discernible. One alternative to PCA is non-negative matrix factorization, but classically this requires the data to also be non-negative, an assumption obviously unmet by neural field time series. However, formulations of nonnegative matrix factorization have been proposed which relax this constraint (Wu & Wang, 2014), and are a promising approach for future studies.

We analyzed human MEG data at the sensor, rather than source level. We therefore inverted the polarity of bursts with positive negative deflections, since they could have originated from unknown source locations and dipole orientations. However source level analyses also suffer from sign ambiguity, requiring laminar LFP data to resolve the true directionality of intracellular currents. Efforts have been made to resolve this inherent source level polarity ambiguity (Rossi & Van Schependom, 2022; Vidaurre et al., 2016), and cortical column estimation in MEG source reconstruction (Bonaiuto et al., 2020) coupled with biophysical modeling can distinguish between competing models (Bonaiuto et al., 2021). However, this is also a strength of this study, as we have shown that beta burst waveform motifs can be identified and classified without source reconstruction, which is especially promising for EEG studies.

Finally, we were unable to account for the most task-modulated waveform motifs by varying the input parameters of the biophysical model. This model was developed to account for beta bursts in somatosensory cortex (S. R. Jones et al., 2009; Law et al., 2022; Sherman et al., 2016), and may require modifications to account for bursts generated in motor cortex. The somatosensory cortex receives its primary inputs from the thalamus (E. G. Jones, 1998, 2001; Mo & Sherman, 2019; W. Zhang & Bruno, 2019), whereas the primary motor cortex has been recently shown to receive strong projections to deep and superficial layers from overlapping neural populations within a wide network of other cortical and subcortical regions including sensory and premotor cortices and the thalamus (Geng et al., 2022). These projections likely contribute to different pre- and post- movement computational processes such as movement planning and evaluation, and the nonlinear combination of these overlapping synaptic drives may thus account for the variable rate-varying burst shapes that we observe.

In the field of neurodegenerative diseases, especially Parkinson’s disease (PD), much effort has been devoted to finding individualized biomarkers that can inform appropriate early interventions (D. B. Miller & O’Callaghan, 2015; Titova & Chaudhuri, 2017). Activity in the beta band, bursts in particular, is instrumentally linked to PD symptomatology (e.g. Little & Brown, 2014; McCarthy et al., 2011). Various burst metrics are related to motor impairments (e.g. Anidi et al., 2018; Kehnemouyi et al., 2021; Lofredi et al., 2019), response to medication (e.g. Duchet et al., 2021; Jackson et al., 2019; Tinkhauser, Pogosyan, Tan, et al., 2017; Yeh et al., 2020), and effects of deep brain stimulation (e.g. Pauls et al., 2022; Schmidt et al., 2020; Tinkhauser, Pogosyan, Little, et al., 2017). All of these metrics were either derived from TF decompositions or band-pass filtered signal amplitude envelopes, and we have shown that TF-based burst features do not differentiate the underlying waveform in the temporal domain. However, the underlying waveform motifs and motif-specific burst rate modulations we observe could offer much needed precision in determining PD biomarkers. Moreover, the sensitivity of such measures for early diagnosis is likely to be far greater than coarser TF-based burst features. Specific waveform motifs could be rapidly detected and targeted with deep brain stimulation devices to deliver more temporally precise interventions. Template matching of waveform motifs in the time domain could potentially reduce the lag between burst detection and stimulation, thus increasing treatment efficacy.

Studies of sensorimotor activity in developing populations typically focus on the alpha or mu frequency bands, and little is known about the development of beta band activity in infancy (Cuevas et al., 2014; Perone & Gartstein, 2019). It has recently been shown that, similar to alpha, there are age-related changes in beta frequency and power (He et al., 2019; Johnson et al., 2019; Rayson et al., 2022; Trevarrow et al., 2019) from infancy to adulthood. In infancy, the peak beta frequency is 15 Hz (Rayson et al., 2022), but movement-related artifacts from facial and arm movements also occur around this frequency (Georgieva et al., 2020). This makes it extremely difficult to separate movement-related neural activity from artifactual activity using TF-based analyses. However, the approach we present could disentangle the two signals. Our adaptive single trial burst detection algorithm detects all potential bursts across a wide range of amplitudes, thus not neglecting bursts of neural activity that are potentially lower in amplitude than movement-related artifacts. Analysis of burst waveform motifs could then separate the cortically generated beta bursts from waveforms corresponding to muscle artifacts even though they overlap in frequency range. Changes in beta power, peak frequency, burst rate and mean waveform have also been found during healthy aging in adulthood (Brady & Bardouille, 2022; Heinrichs-Graham et al., 2018; Heinrichs-Graham & Wilson, 2016; Rempe et al., 2022; Rossiter et al., 2014). Future studies applying our approach to such data could help provide mechanistic explanations for these age-related changes at the individual level.

The vast majority of non-invasive brain-computer interfaces (BCIs) try to identify and characterize single imagined movements using temporally averaged power in the mu and beta bands (Brodu et al., 2011; Herman et al., 2008; Pfurtscheller & Neuper, 2001). Most recent advances in the field rely on sophisticated machine learning techniques (Barachant et al., 2012; Llera et al., 2014; Lotte et al., 2018; Song et al., 2013), but we have recently argued that what is needed for further predictive power is a more fine-grained approach to feature extraction aimed at the temporal signatures of bursts of mu and beta activity (Papadopoulos et al., 2022). We have here demonstrated that averaged beta power includes many burst motifs that do not change in rate pre-movement. By filtering out these events and focusing on task-modulated burst motifs, the SNR of the features fed into machine learning algorithms for BCI could be greatly increased, improving classification accuracy. Rather than PCA, supervised or semi-supervised dimensionality reduction techniques such as demixed PCA (Kobak et al., 2016) or a common spatial pattern approach in the time domain (CSP; Congedo et al., 2016) could be used to determine burst waveform motifs whose rate modulations maximally distinguish between movement types. Finally, given a training dataset to define burst motifs and their modulations by the task, an online algorithm could be developed using template matching to detect bursts with particular waveforms.

Whilst TF decomposition has proven useful for segregating and identifying classes of frequency-specific neural activity, TF-based features that do not include phase information are ambiguous with respect to the underlying temporal waveform shape. Our results demonstrate that this information is crucial for determining which bursts drive movement-related changes in beta activity. Sensorimotor beta activity can therefore be decomposed into distinct burst types which differ in their rate-based dynamics, and likely index different computational processes. This is unlikely to be unique to the beta frequency band, and thus underscores the importance and power of analyzing frequency-specific neural activity in the temporal domain.

## Acknowledgements

This research was supported by grants from the European Research Council (ERC) under the European Union’s Horizon 2020 research and innovation programme (ERC consolidator grant 864550; ERC starting grant 716862), and the French National Research Agency (ANR) within the program “Investissements d’Avenir” (2019-ANR-LABX-02). The funders had no role in the preparation of the manuscript. We thank F Lamberton for assistance with MRI sequence development.

## Supplementary Figures

**Figure S1.**
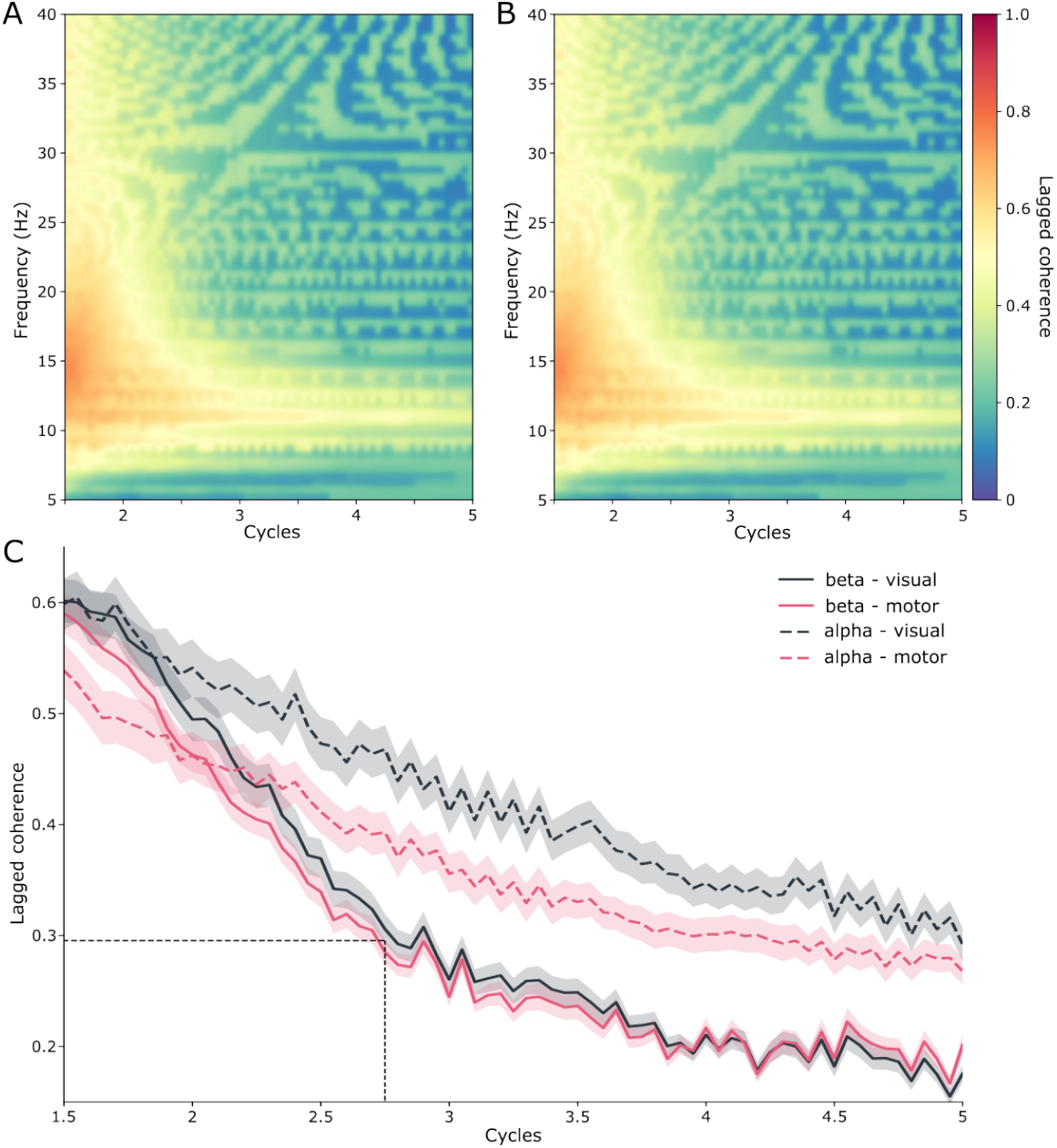
(A) Mean lagged coherence over subjects during the visual epochs from 5 to 40 Hz and 1.5 to 5 cycles. Lagged coherence is high in the alpha and beta bands from 1.5 to 2 cycles, but then drops off more quickly in the beta band with increasing cycles. (B) As in A during the motor epochs. (C) Lagged coherence during the visual (black lines) and motor (red lines) averaged over the alpha (dashed lines) and beta (solid lines) frequency bands. Half of the FWHM (dotted lines) was used to define the width of the time window for burst waveform extraction.

**Figure S2.**
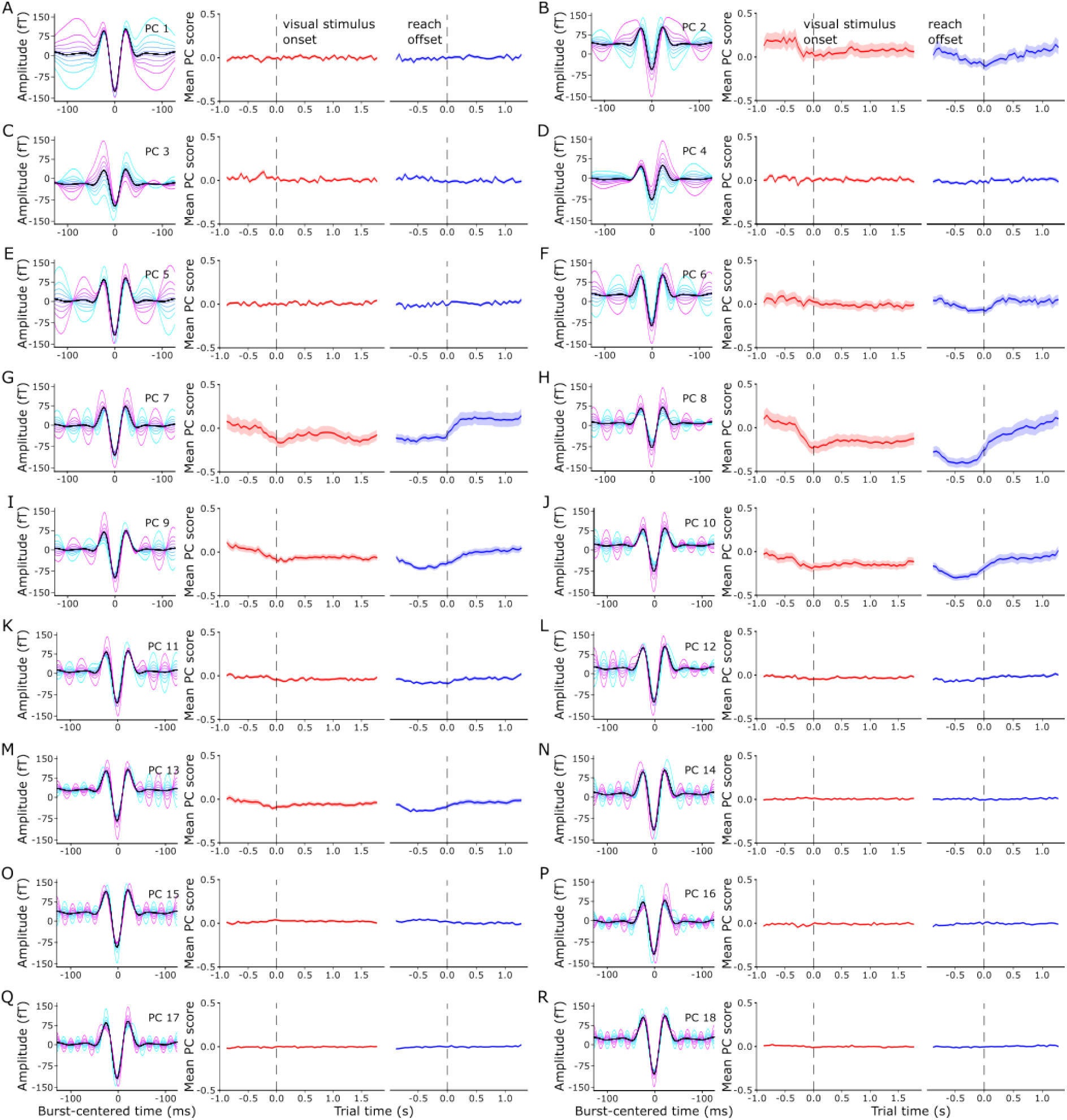
(A-R) For each significant component, mean burst waveform shapes of bursts with a score within the 0 - 10th (cyan) to 90 - 100th (magenta) percentile of all burst scores for that component (left panels). The panels on the right show the time course of the mean burst score (solid line; shaded region shows SEM) for each component during the visual and motor epochs.

**Figure S3.**
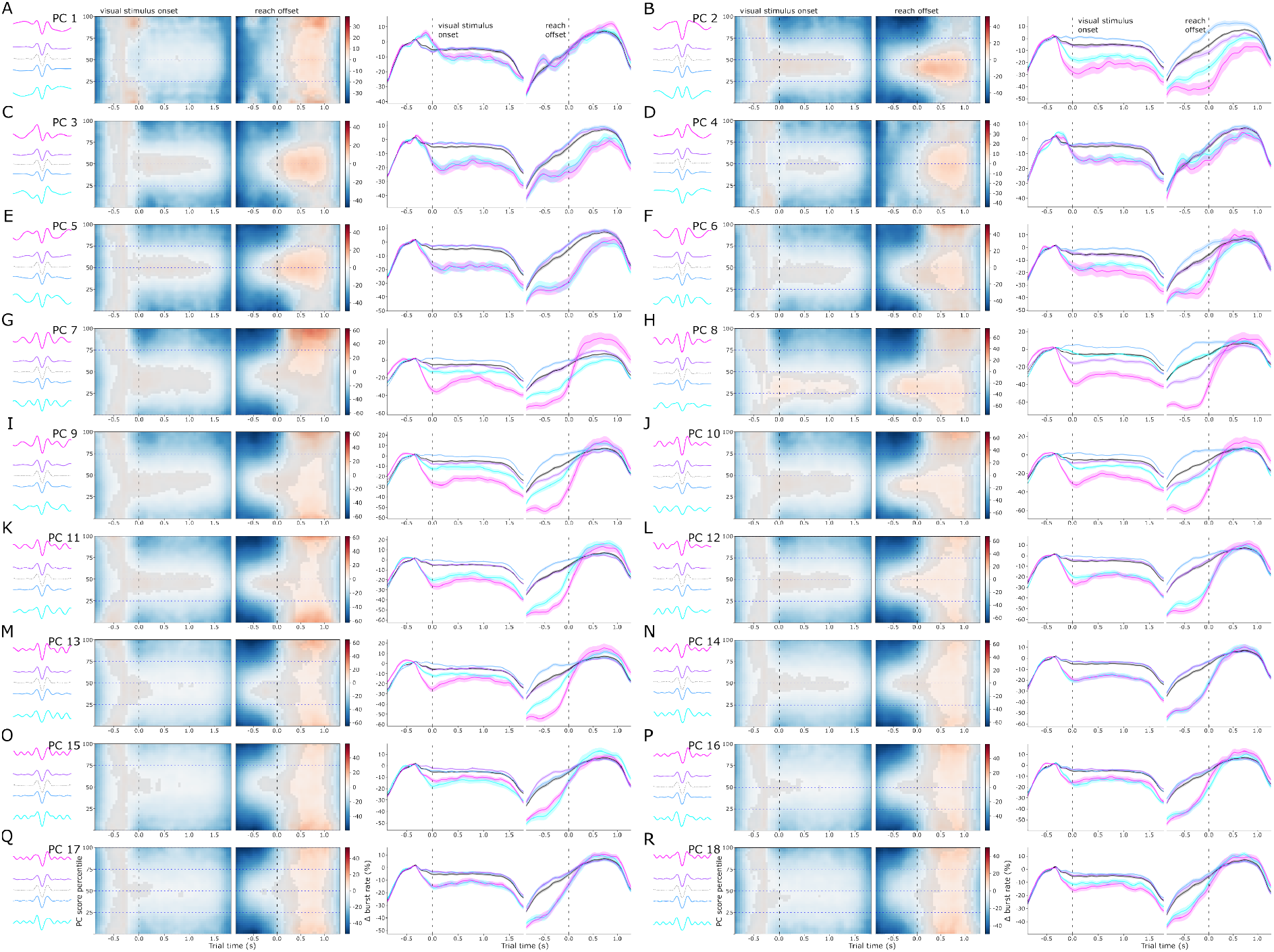
(A-R) The mean waveforms of bursts with scores in each quartile of the range of scores for PCs 1-18 (left panels, colored lines) and the mean overall burst waveform (black), and the mean burst rate over time and across the range of component scores during the visual and motor epochs (middle panels). The gray overlays obscure the time points and score ranges where the rate did not differ from the baseline rate. The horizontal dashed lines indicate the quartile limits. The mean baseline-corrected rate of bursts with scores in each quartile of the range of scores along each component (right panels, colored lines, the shaded area indicates the SEM) over the course of the visual and motor epochs. The black line represents the mean rate of all bursts (the shaded area indicates the SEM).

**Figure S4.**
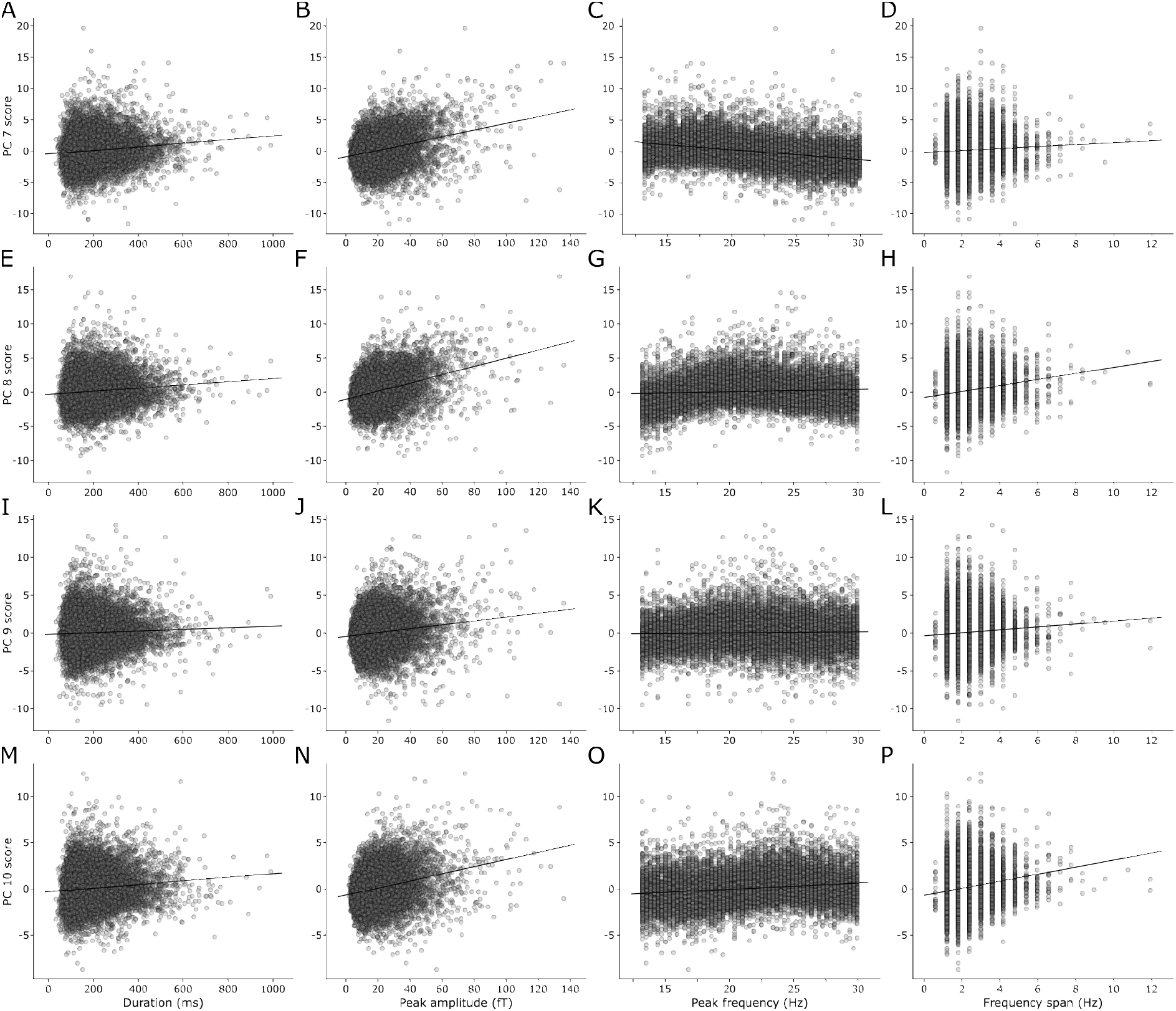
(A-D) The relationship between burst duration (A), peak amplitude (B), peak frequency (C), and frequency span (D) with PC 7 score. (E-P) As in A-D for PC 8 (E-H), PC 9 (I-L), and PC 10 (M-P).

**Figure S5.**
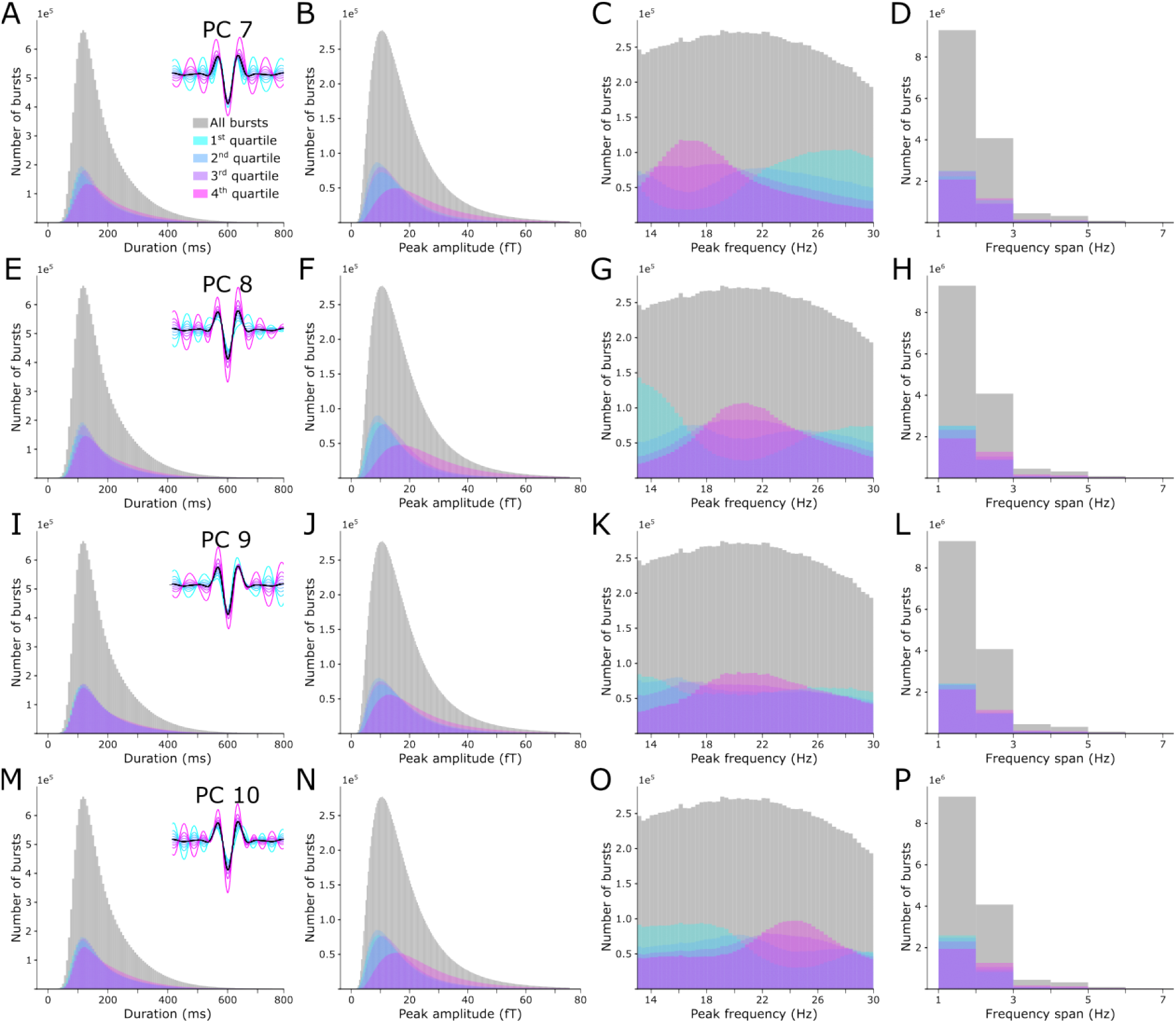
The distributions of burst duration, peak amplitude, peak frequency, and frequency span for all bursts, and separately for bursts with scores in the 1st - 4th percentile of components 7 (A-D), 8 (E-H), 9 (I-L), and 10 (M-P). The gray histograms show the distribution of each feature for all bursts, and the colored histograms represent the distributions of bursts with scores from the 1st (cyan) to 4th (magenta) quartile for each component.

**Figure S6.**
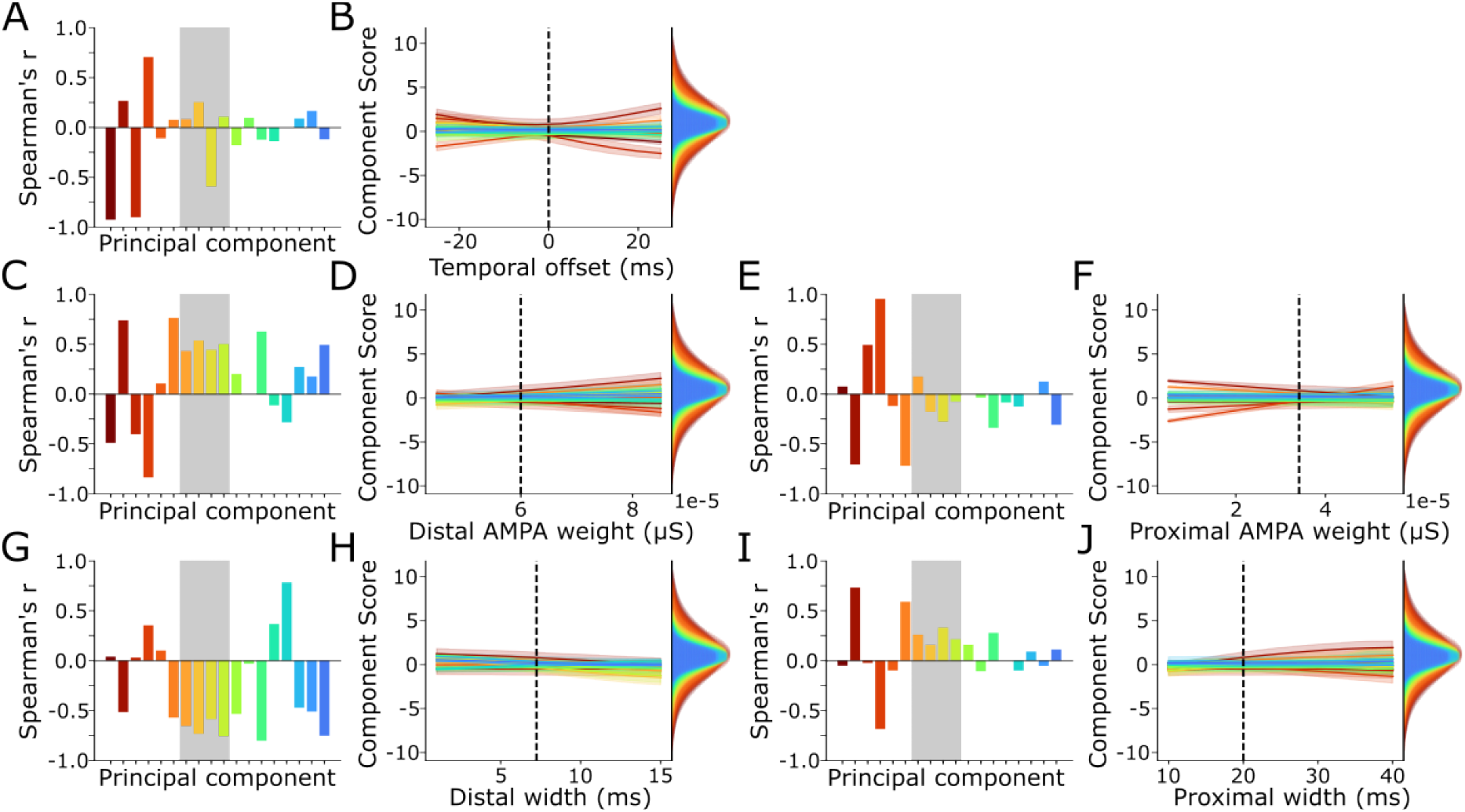
(A) The correlation (Spearman’s ρ) between temporal offset of the proximal and distal drives and the score of the simulated burst waveforms for each PC. The gray rectangle highlights the selected task-modulated components (7-10). (B) The mean score of the simulated bursts at each level of temporal offset tested (shaded area shows SD), for each PC. The vertical dashed line indicates the value used in the base simulations. The histograms show the distribution of scores for each PC for bursts from the human MEG data. (C-J) As in (A-B) for distal drive strength (C-D), proximal drive strength (E-F), distal drive width (G-H), and proximal drive width (I-J).

**Table S1.**
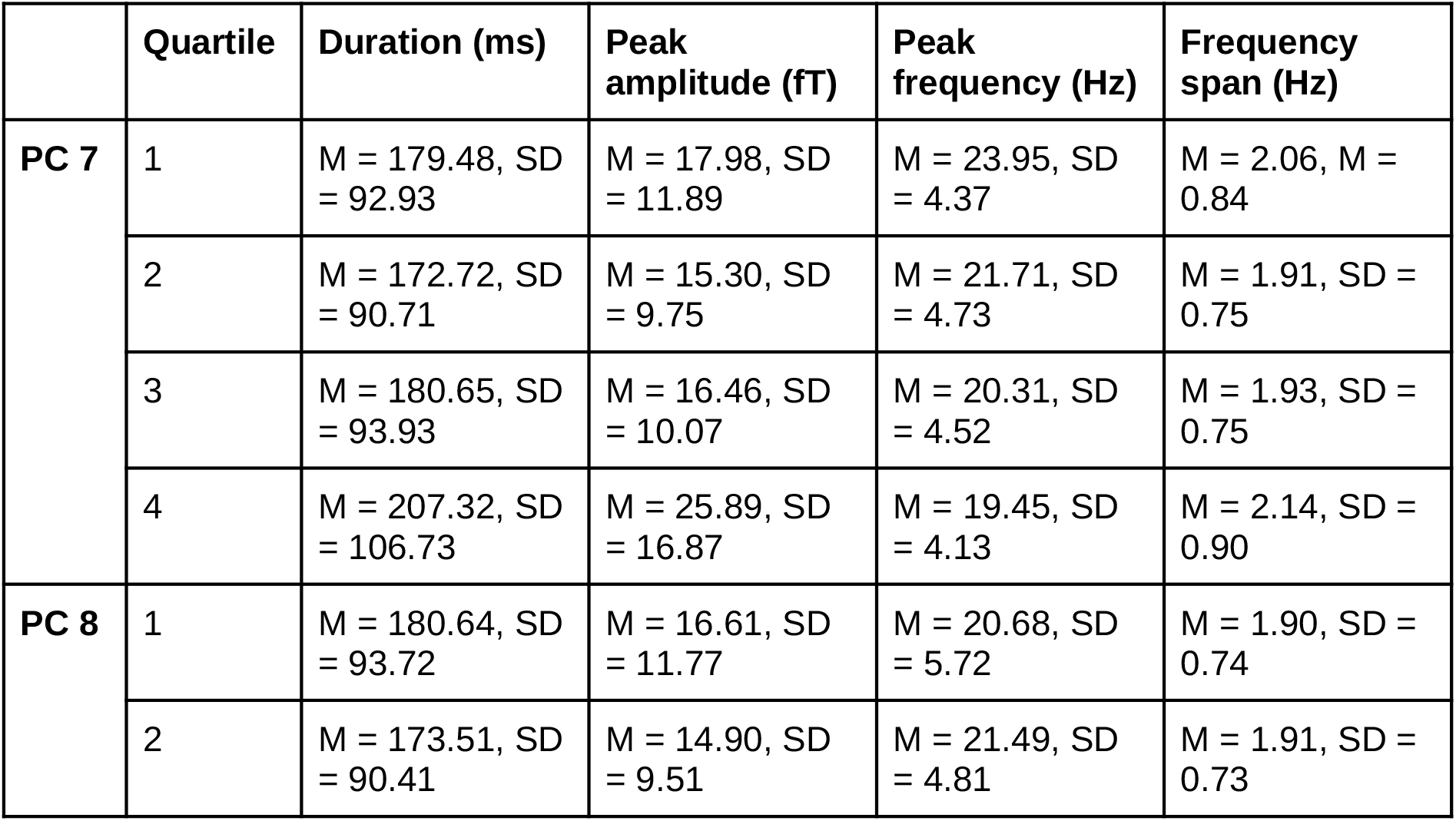
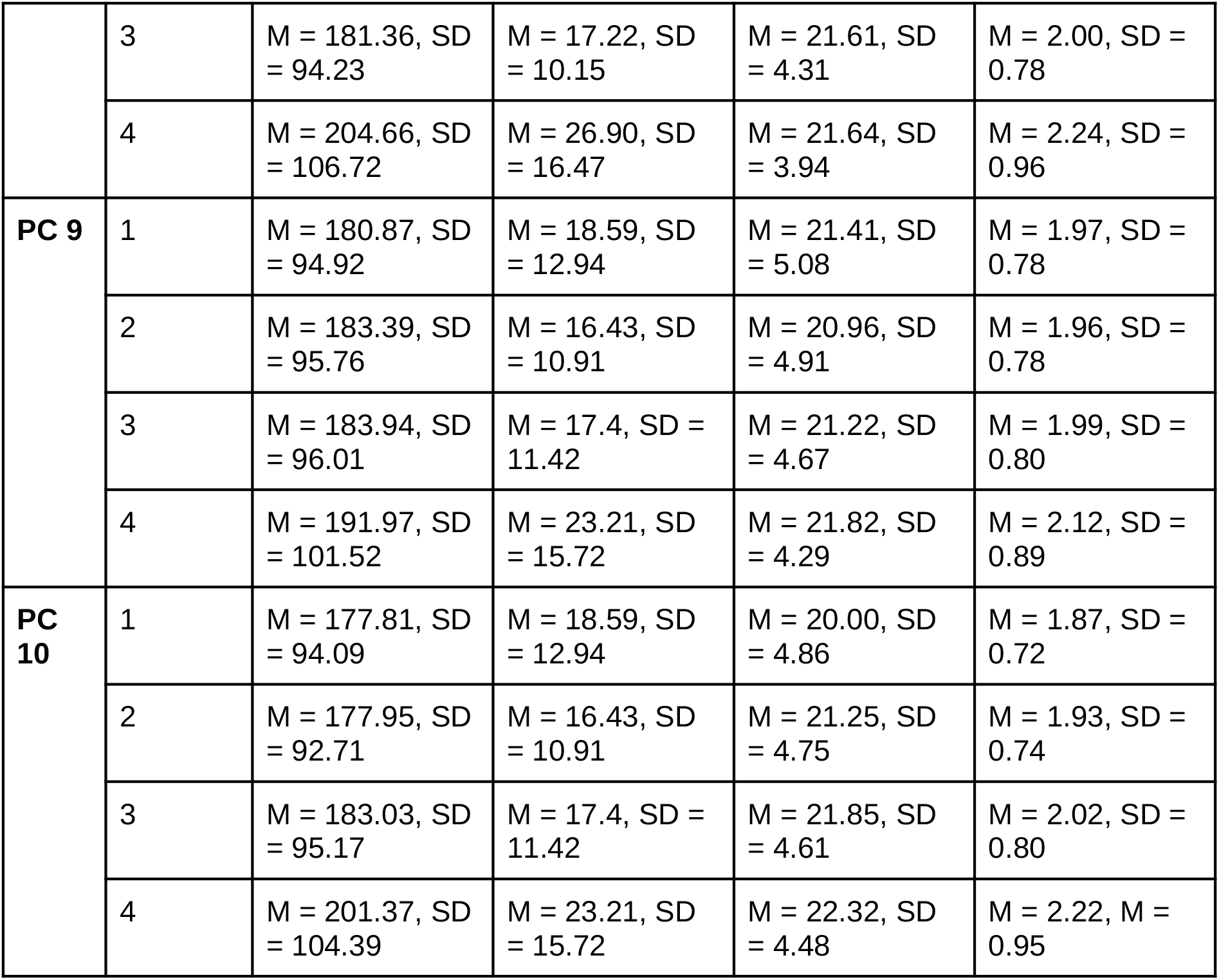
The mean and standard deviation of burst duration, peak amplitude, peak frequency, and frequency span for bursts with scores in the 1st - 4th percentile of components 7, 8, 9, and 10.

|  | Quartile | Duration (ms) | Peak amplitude (fT) | Peak frequency (Hz) | Frequency span (Hz) |
| --- | --- | --- | --- | --- | --- |
| **PC 7** | 1 | M = 179.48, SD = 92.93 | M = 17.98, SD = 11.89 | M = 23.95, SD = 4.37 | M = 2.06, M = 0.84 |
|  | 2 | M = 172.72, SD = 90.71 | M = 15.30, SD = 9.75 | M = 21.71, SD = 4.73 | M = 1.91, SD = 0.75 |
|  | 3 | M = 180.65, SD = 93.93 | M = 16.46, SD = 10.07 | M = 20.31, SD = 4.52 | M = 1.93, SD = 0.75 |
|  | 4 | M = 207.32, SD = 106.73 | M = 25.89, SD = 16.87 | M = 19.45, SD = 4.13 | M = 2.14, SD = 0.90 |
| **PC 8** | 1 | M = 180.64, SD = 93.72 | M = 16.61, SD = 11.77 | M = 20.68, SD = 5.72 | M = 1.90, SD = 0.74 |
|  | 2 | M = 173.51, SD = 90.41 | M = 14.90, SD = 9.51 | M = 21.49, SD = 4.81 | M = 1.91, SD = 0.73 |
|  | 3 | M = 181.36, SD = 94.23 | M = 17.22, SD = 10.15 | M = 21.61, SD = 4.31 | M = 2.00, SD = 0.78 |
|  | 4 | M = 204.66, SD = 106.72 | M = 26.90, SD = 16.47 | M = 21.64, SD = 3.94 | M = 2.24, SD = 0.96 |
| **PC 9** | 1 | M = 180.87, SD = 94.92 | M = 18.59, SD = 12.94 | M = 21.41, SD = 5.08 | M = 1.97, SD = 0.78 |
|  | 2 | M = 183.39, SD = 95.76 | M = 16.43, SD = 10.91 | M = 20.96, SD = 4.91 | M = 1.96, SD = 0.78 |
|  | 3 | M = 183.94, SD = 96.01 | M = 17.4, SD = 11.42 | M = 21.22, SD = 4.67 | M = 1.99, SD = 0.80 |
|  | 4 | M = 191.97, SD = 101.52 | M = 23.21, SD = 15.72 | M = 21.82, SD = 4.29 | M = 2.12, SD = 0.89 |
| **PC 10** | 1 | M = 177.81, SD = 94.09 | M = 18.59, SD = 12.94 | M = 20.00, SD = 4.86 | M = 1.87, SD = 0.72 |
|  | 2 | M = 177.95, SD = 92.71 | M = 16.43, SD = 10.91 | M = 21.25, SD = 4.75 | M = 1.93, SD = 0.74 |
|  | 3 | M = 183.03, SD = 95.17 | M = 17.4, SD = 11.42 | M = 21.85, SD = 4.61 | M = 2.02, SD = 0.80 |
|  | 4 | M = 201.37, SD = 104.39 | M = 23.21, SD = 15.72 | M = 22.32, SD = 4.48 | M = 2.22, SD = 0.95 |

**Table S2.** Pairwise, Bonferonni-corrected comparisons between the quartiles of PCs 7, 8, 9, and 10 in terms of the distribution of burst duration.

|  | 1st - 2nd | 1st - 3rd | 1st - 4th | 2nd - 3rd | 2nd - 4th | 3rd - 4th |
| --- | --- | --- | --- | --- | --- | --- |
| **PC 7** | D = 0.050, p < 0.001 | D = 0.030, p < 0.001 | D = 0.127, p < 0.001 | D = 0.056, p < 0.001 | D = 0.177, p < 0.001 | D = 0.134, p < 0.001 |
| **PC 8** | D = 0.048, p < 0.001 | D = 0.036, p < 0.001 | D = 0.086, p < 0.001 | D = 0.044, p < 0.001 | D = 0.143, p < 0.001 | D = 0.108, p < 0.001 |
| **PC 9** | D = 0.022, p < 0.001 | D = 0.026, p < 0.001 | D = 0.042, p < 0.001 | D = 0.005, p < 0.001 | D = 0.037, p < 0.001 | D = 0.037, p < 0.001 |
| **PC 10** | D = 0.023, p < 0.001 | D = 0.038, p < 0.001 | D = 0.107, p < 0.001 | D = 0.025, p < 0.001 | D = 0.115, p < 0.001 | D = 0.092, p < 0.001 |

**Table S3.** Pairwise, Bonferonni-corrected comparisons between the quartiles of PCs 7, 8, 9, and 10 in terms of the distribution of burst peak amplitude.

|  | 1st - 2nd | 1st - 3rd | 1st - 4th | 2nd - 3rd | 2nd - 4th | 3rd - 4th |
| --- | --- | --- | --- | --- | --- | --- |
| **PC 7** | D = 0.108,<br>p < 0.001 | D = 0.054,<br>p < 0.001 | D = 0.242,<br>p < 0.001 | D = 0.064,<br>p < 0.001 | D = 0.344,<br>p < 0.001 | D = 0.296,<br>p < 0.001 |
| **PC 8** | D = 0.059,<br>p < 0.001 | D = 0.080,<br>p < 0.001 | D = 0.351,<br>p < 0.001 | D = 0.126,<br>p < 0.001 | D = 0.411,<br>p < 0.001 | D = 0.313,<br>p < 0.001 |
| **PC 9** | D = 0.075,<br>p < 0.001 | D = 0.036,<br>p < 0.001 | D = 0.148,<br>p < 0.001 | D = 0.040,<br>p < 0.001 | D = 0.222,<br>p < 0.001 | D = 0.184,<br>p < 0.001 |
| **PC 10** | D = 0.053,<br>p < 0.001 | D = 0.045,<br>p < 0.001 | D = 0.274,<br>p < 0.001 | D = 0.085,<br>p < 0.001 | D = 0.326,<br>p < 0.001 | D = 0.255,<br>p < 0.001 |

**Table S4.** Pairwise, Bonferonni-corrected comparisons between the quartiles of PCs 7, 8, 9, and 10 in terms of the distribution of burst peak frequency.

|  | **1st - 2nd** | **1st - 3rd** | **1st - 4th** | **2nd - 3rd** | **2nd - 4th** | **3rd - 4th** |
| --- | --- | --- | --- | --- | --- | --- |
| **PC 7** | D = 0.227,<br>p < 0.001 | D = 0.374,<br>p < 0.001 | D = 0.472,<br>p < 0.001 | D = 0.156,<br>p < 0.001 | D = 0.261,<br>p < 0.001 | D = 0.107,<br>p < 0.001 |
| **PC 8** | D = 0.173,<br>p < 0.001 | D = 0.238,<br>p < 0.001 | D = 0.278,<br>p < 0.001 | D = 0.070,<br>p < 0.001 | D = 0.117,<br>p < 0.001 | D = 0.048,<br>p < 0.001 |
| **PC 9** | D = 0.050,<br>p < 0.001 | D = 0.049,<br>p < 0.001 | D = 0.113,<br>p < 0.001 | D = 0.047,<br>p < 0.001 | D = 0.135,<br>p < 0.001 | D = 0.092,<br>p < 0.001 |
| **PC 10** | D = 0.153,<br>p < 0.001 | D = 0.232,<br>p < 0.001 | D = 0.294,<br>p < 0.001 | D = 0.085,<br>p < 0.001 | D = 0.153,<br>p < 0.001 | D = 0.070,<br>p < 0.001 |

**Table S5.** Pairwise, Bonferonni-corrected comparisons between the quartiles of PCs 7, 8, 9, and 10 in terms of the distribution of frequency span.

|  | **1st - 2nd** | **1st - 3rd** | **1st - 4th** | **2nd - 3rd** | **2nd - 4th** | **3rd - 4th** |
| --- | --- | --- | --- | --- | --- | --- |
| **PC 7** | D = 0.078,<br>p < 0.001 | D = 0.069,<br>p < 0.001 | D = 0.042,<br>p < 0.001 | D = 0.016,<br>p < 0.001 | D = 0.120,<br>p < 0.001 | D = 0.111,<br>p < 0.001 |
| **PC 8** | D = 0.017,<br>p < 0.001 | D = 0.063,<br>p < 0.001 | D = 0.174,<br>p < 0.001 | D = 0.055,<br>p < 0.001 | D = 0.171,<br>p < 0.001 | D = 0.115,<br>p < 0.001 |
| **PC 9** | D = 0.004,<br>p < 0.001 | D = 0.013,<br>p < 0.001 | D = 0.076,<br>p < 0.001 | D = 0.017,<br>p < 0.001 | D = 0.081,<br>p < 0.001 | D = 0.064,<br>p < 0.001 |
| **PC 10** | D = 0.044,<br>p < 0.001 | D = 0.084,<br>p < 0.001 | D = 0.184,<br>p < 0.001 | D = 0.050,<br>p < 0.001 | D = 0.150,<br>p < 0.001 | D = 0.100,<br>p < 0.001 |

## References

Alayrangues, J., Torrecillos, F., Jahani, A., & Malfait, N. (2019). Error-related modulations of the sensorimotor post-movement and foreperiod beta-band activities arise from distinct neural substrates and do not reflect efferent signal processing. NeuroImage, 184, 10–24. https://doi.org/10.1016/j.neuroimage.2018.09.013

Anidi, C., O’Day, J. J., Anderson, R. W., Afzal, M. F., Syrkin-Nikolau, J., Velisar, A., & Bronte-Stewart, H. M. (2018). Neuromodulation targets pathological not physiological beta bursts during gait in Parkinson’s disease. Neurobiology of Disease, 120, 107–117. https://doi.org/10.1016/j.nbd.2018.09.004

Baker, S. N., Olivier, E., & Lemon, R. N. (1997). Coherent oscillations in monkey motor cortex and hand muscle EMG show task-dependent modulation. The Journal of Physiology, 501(1), 225–241. https://doi.org/10.1111/j.1469-7793.1997.225bo.x

Barachant, A., Bonnet, S., Congedo, M., & Jutten, C. (2012). Multiclass brain-computer interface classification by Riemannian geometry. IEEE Transactions on Bio-Medical Engineering, 59(4), 920–928. https://doi.org/10.1109/TBME.2011.2172210

Bartolo, R., & Merchant, H. (2015). β Oscillations Are Linked to the Initiation of Sensory-Cued Movement Sequences and the Internal Guidance of Regular Tapping in the Monkey. Journal of Neuroscience, 35(11), 4635–4640. https://doi.org/10.1523/JNEUROSCI.4570-14.2015

Bates, D., Mächler, M., Bolker, B., & Walker, S. (2014). Fitting linear mixed-effects models using lme4. ArXiv Preprint ArXiv:1406.5823.

Bonaiuto, J. J., Afdideh, F., Ferez, M., Wagstyl, K., Mattout, J., Bonnefond, M., Barnes, G. R., & Bestmann, S. (2020). Estimates of cortical column orientation improve MEG source inversion. NeuroImage, 216, 116862. https://doi.org/10.1016/j.neuroimage.2020.116862

Bonaiuto, J. J., Little, S., Neymotin, S. A., Jones, S. R., Barnes, G. R., & Bestmann, S. (2021). Laminar dynamics of high amplitude beta bursts in human motor cortex. NeuroImage, 242, 118479. https://doi.org/10.1016/j.neuroimage.2021.118479

Bonaiuto, J. J., Rossiter, H. E., Meyer, S. S., Adams, N., Little, S., Callaghan, M. F., Dick, F., Bestmann, S., & Barnes, G. R. (2018). Non-invasive laminar inference with MEG: Comparison of methods and source inversion algorithms. NeuroImage, 167, 372–383. https://doi.org/10.1016/j.neuroimage.2017.11.068

Boonstra, T. W., Daffertshofer, A., Breakspear, M., & Beek, P. J. (2007). Multivariate time– frequency analysis of electromagnetic brain activity during bimanual motor learning. NeuroImage, 36(2), 370–377. https://doi.org/10.1016/j.neuroimage.2007.03.012

Boto, E., Shah, V., Hill, R. M., Rhodes, N., Osborne, J., Doyle, C., Holmes, N., Rea, M., Leggett, J., Bowtell, R., & Brookes, M. J. (2022). Triaxial detection of the neuromagnetic field using optically-pumped magnetometry: Feasibility and application in children. NeuroImage, 252, 119027. https://doi.org/10.1016/j.neuroimage.2022.119027

Brady, B., & Bardouille, T. (2022). Periodic/Aperiodic parameterization of transient oscillations (PAPTO)–Implications for healthy ageing. NeuroImage, 251, 118974. https://doi.org/10.1016/j.neuroimage.2022.118974

Brainard, D. H. (1997). The psychophysics toolbox. Spatial Vision, 10(4), 433–436.

Brodu, N., Lotte, F., & Lécuyer, A. (2011). Comparative Study of Band-Power Extraction Techniques for Motor Imagery Classification. 1. https://doi.org/10.1109/CCMB.2011.5952105

Cagnan, H., Mallet, N., Moll, C. K. E., Gulberti, A., Holt, A. B., Westphal, M., Gerloff, C., Engel, A. K., Hamel, W., Magill, P. J., Brown, P., & Sharott, A. (2019). Temporal evolution of beta bursts in the parkinsonian cortical and basal ganglia network. Proceedings of the National Academy of Sciences, 116(32), 16095–16104. https://doi.org/10.1073/pnas.1819975116

Cassim, F., Monaca, C., Szurhaj, W., Bourriez, J.-L., Defebvre, L., Derambure, P., & Guieu, J.-D. (2001). Does post-movement beta synchronization reflect an idling motor cortex? Neuroreport, 12(17), 3859–3863.

Cheyne, D. O. (2013). MEG studies of sensorimotor rhythms: A review. Experimental Neurology, 245, 27–39. https://doi.org/10.1016/j.expneurol.2012.08.030

Cole, S., Meij, R. van der, Peterson, E. J., Hemptinne, C. de, Starr, P. A., & Voytek, B. (2017). Nonsinusoidal Beta Oscillations Reflect Cortical Pathophysiology in Parkinson’s Disease. Journal of Neuroscience, 37(18), 4830–4840. https://doi.org/10.1523/JNEUROSCI.2208-16.2017

Cole, S., & Voytek, B. (2017). Brain Oscillations and the Importance of Waveform Shape. Trends in Cognitive Sciences, 21(2), 137–149. https://doi.org/10.1016/j.tics.2016.12.008

Cole, S., & Voytek, B. (2019). Cycle-by-cycle analysis of neural oscillations. Journal of Neurophysiology, 122(2), 849–861. https://doi.org/10.1152/jn.00273.2019

Congedo, M., Korczowski, L., Delorme, A., & Lopes da silva, F. (2016). Spatio-temporal common pattern: A companion method for ERP analysis in the time domain. Journal of Neuroscience Methods, 267, 74–88. https://doi.org/10.1016/j.jneumeth.2016.04.008

Cuevas, K., Cannon, E. N., Yoo, K., & Fox, N. A. (2014). The infant EEG mu rhythm: Methodological considerations and best practices. Developmental Review, 34(1), 26–43. https://doi.org/10.1016/j.dr.2013.12.001

de Cheveigné, A. (2020). ZapLine: A simple and effective method to remove power line artifacts. NeuroImage, 207, 116356. https://doi.org/10.1016/j.neuroimage.2019.116356

Diesburg, D. A., Greenlee, J. D., & Wessel, J. R. (2021). Cortico-subcortical β burst dynamics underlying movement cancellation in humans. ELife, 10, e70270. https://doi.org/10.7554/eLife.70270

Donner, T. H., Siegel, M., Fries, P., & Engel, A. K. (2009). Buildup of Choice-Predictive Activity in Human Motor Cortex during Perceptual Decision Making. Current Biology, 19(18), 1581–1585. https://doi.org/10.1016/j.cub.2009.07.066

Donoghue, T., Haller, M., Peterson, E. J., Varma, P., Sebastian, P., Gao, R., Noto, T., Lara, A. H., Wallis, J. D., Knight, R. T., Shestyuk, A., & Voytek, B. (2020). Parameterizing neural power spectra into periodic and aperiodic components. Nature Neuroscience, 23(12), Article 12. https://doi.org/10.1038/s41593-020-00744-x

Duchet, B., Ghezzi, F., Weerasinghe, G., Tinkhauser, G., Kühn, A. A., Brown, P., Bick, C., & Bogacz, R. (2021). Average beta burst duration profiles provide a signature of dynamical changes between the ON and OFF medication states in Parkinson’s disease. PLoS Computational Biology, 17(7), e1009116. https://doi.org/10.1371/journal.pcbi.1009116

Echeverria-Altuna, I., Quinn, A. J., Zokaei, N., Woolrich, M. W., Nobre, A. C., & van Ede, F. (2021). Transient beta activity and connectivity during sustained motor behaviour [Preprint]. Neuroscience. https://doi.org/10.1101/2021.03.02.433514

Engel, A. K., & Fries, P. (2010). Beta-band oscillations-signalling the status quo? Current Opinion in Neurobiology, 20(2), 156–165. https://doi.org/10.1016/j.conb.2010.02.015

Enz, N., Ruddy, K. L., Rueda-Delgado, L. M., & Whelan, R. (2021). Volume of β-Bursts, But Not Their Rate, Predicts Successful Response Inhibition. Journal of Neuroscience, 41(23), 5069–5079. https://doi.org/10.1523/JNEUROSCI.2231-20.2021

Erbil, N., & Ungan, P. (2007). Changes in the alpha and beta amplitudes of the central EEG during the onset, continuation, and offset of long-duration repetitive hand movements. Brain Research, 1169, 44–56. https://doi.org/10.1016/j.brainres.2007.07.014

Feingold, J., Gibson, D. J., DePasquale, B., & Graybiel, A. M. (2015). Bursts of beta oscillation differentiate postperformance activity in the striatum and motor cortex of monkeys performing movement tasks. Proceedings of the National Academy of Sciences, 112(44), 13687–13692. https://doi.org/10.1073/pnas.1517629112

Fine, J. M., Moore, D., & Santello, M. (2017). Neural oscillations reflect latent learning states underlying dual-context sensorimotor adaptation. NeuroImage, 163, 93–105. https://doi.org/10.1016/j.neuroimage.2017.09.026

Fischl, B., Salat, D. H., Busa, E., Albert, M., Dieterich, M., Haselgrove, C., van der Kouwe, A., Killiany, R., Kennedy, D., Klaveness, S., Montillo, A., Makris, N., Rosen, B., & Dale, A. M. (2002). Whole Brain Segmentation: Automated Labeling of Neuroanatomical Structures in the Human Brain. Neuron, 33(3), 341–355. https://doi.org/10.1016/S0896-6273(02)00569-X

Fox, J., Weisberg, S., Price, B., Adler, D., Bates, D., Baud-Bovy, G., & Bolker, B. (2019). car: Companion to Applied Regression. R package version 3.0-2. Website Https://CRAN.R-Project.Org/Package=Car [Accessed 17 March 2020].

Fransen, A. M. M., van Ede, F., & Maris, E. (2015). Identifying neuronal oscillations using rhythmicity. NeuroImage, 118, 256–267. https://doi.org/10.1016/j.neuroimage.2015.06.003

Geng, H.-Y., Arbuthnott, G., Yung, W.-H., & Ke, Y. (2022). Long-range monosynaptic inputs targeting apical and basal dendrites of primary motor cortex deep output neurons. Cerebral Cortex, 32(18), 3975–3989. https://doi.org/10.1093/cercor/bhab460

Georgieva, S., Lester, S., Noreika, V., Yilmaz, M. N., Wass, S., & Leong, V. (2020). Toward the Understanding of Topographical and Spectral Signatures of Infant Movement Artifacts in Naturalistic EEG. Frontiers in Neuroscience, 14. https://www.frontiersin.org/articles/10.3389/fnins.2020.00352

Gramfort, A., Luessi, M., Larson, E., Engemann, D. A., Strohmeier, D., Brodbeck, C., Parkkonen, L., & Hämäläinen, M. S. (2014). MNE software for processing MEG and EEG data. NeuroImage, 86, 446–460. https://doi.org/10.1016/j.neuroimage.2013.10.027

Haegens, S., Nácher, V., Hernández, A., Luna, R., Jensen, O., & Romo, R. (2011). Beta oscillations in the monkey sensorimotor network reflect somatosensory decision making. Proceedings of the National Academy of Sciences, 108(26), 10708–10713. https://doi.org/10.1073/pnas.1107297108

Haufler, D., Liran, O., Buchanan, R. J., & Pare, D. (2022). Human anterior insula signals salience and deviations from expectations via bursts of beta oscillations. Journal of Neurophysiology. https://doi.org/10.1152/jn.00106.2022

He, W., Donoghue, T., Sowman, P. F., Seymour, R. A., Brock, J., Crain, S., Voytek, B., & Hillebrand, A. (2019). Co-Increasing Neuronal Noise and Beta Power in the Developing Brain [Preprint]. Neuroscience. https://doi.org/10.1101/839258

Heideman, S. G., Quinn, A. J., Woolrich, M. W., van Ede, F., & Nobre, A. C. (2020). Dissecting beta-state changes during timed movement preparation in Parkinson’s disease. Progress in Neurobiology, 184, 101731. https://doi.org/10.1016/j.pneurobio.2019.101731

Heinrichs-Graham, E., Arpin, D. J., & Wilson, T. W. (2016). Cue-related Temporal Factors Modulate Movement-related Beta Oscillatory Activity in the Human Motor Circuit. Journal of Cognitive Neuroscience, 28(7), 1039–1051. https://doi.org/10.1162/jocn_a_00948

Heinrichs-Graham, E., McDermott, T. J., Mills, M. S., Wiesman, A. I., Wang, Y.-P., Stephen, J. M., Calhoun, V. D., & Wilson, T. W. (2018). The lifespan trajectory of neural oscillatory activity in the motor system. Developmental Cognitive Neuroscience, 30, 159–168. https://doi.org/10.1016/j.dcn.2018.02.013

Heinrichs-Graham, E., & Wilson, T. W. (2016). Is an absolute level of cortical beta suppression required for proper movement? Magnetoencephalographic evidence from healthy aging. NeuroImage, 134, 514–521. https://doi.org/10.1016/j.neuroimage.2016.04.032

Herman, P., Prasad, G., McGinnity, T. M., & Coyle, D. (2008). Comparative analysis of spectral approaches to feature extraction for EEG-based motor imagery classification. IEEE Transactions on Neural Systems and Rehabilitation Engineering: A Publication of the IEEE Engineering in Medicine and Biology Society, 16(4), 317–326. https://doi.org/10.1109/TNSRE.2008.926694

Higgins, C., van Es, M. W. J., Quinn, A. J., Vidaurre, D., & Woolrich, M. W. (2022). The relationship between frequency content and representational dynamics in the decoding of neurophysiological data. NeuroImage, 260, 119462. https://doi.org/10.1016/j.neuroimage.2022.119462

Houweling, S., Daffertshofer, A., van Dijk, B. W., & Beek, P. J. (2008). Neural changes induced by learning a challenging perceptual-motor task. NeuroImage, 41(4), 1395–1407. https://doi.org/10.1016/j.neuroimage.2008.03.023

Howe, M. W., Atallah, H. E., McCool, A., Gibson, D. J., & Graybiel, A. M. (2011). Habit learning is associated with major shifts in frequencies of oscillatory activity and synchronized spike firing in striatum. Proceedings of the National Academy of Sciences, 108(40), 16801–16806. https://doi.org/10.1073/pnas.1113158108

Jackson, N., Cole, S. R., Voytek, B., & Swann, N. C. (2019). Characteristics of Waveform Shape in Parkinson’s Disease Detected with Scalp Electroencephalography. ENeuro, 6(3). https://doi.org/10.1523/ENEURO.0151-19.2019

Jasper, H., & Penfield, W. (1949). Electrocorticograms in man: Effect of voluntary movement upon the electrical activity of the precentral gyrus. Archiv Fur Psychiatrie Und Nervenkrankheiten, 183(1–2), 163–174. https://doi.org/10.1007/BF01062488

Jensen, O., Goel, P., Kopell, N., Pohja, M., Hari, R., & Ermentrout, B. (2005). On the human sensorimotor-cortex beta rhythm: Sources and modeling. NeuroImage, 26(2), 347–355. https://doi.org/10.1016/j.neuroimage.2005.02.008

Johnson, B., Jobst, C., Al-Loos, R., He, W., & Cheyne, D. (2019). Developmental Changes in Movement Related Brain Activity in Early Childhood (p. 531905). bioRxiv. https://doi.org/10.1101/531905

Jones, E. G. (1998). Viewpoint: The core and matrix of thalamic organization. Neuroscience, 85(2), 331–345. https://doi.org/10.1016/S0306-4522(97)00581-2

Jones, E. G. (2001). The thalamic matrix and thalamocortical synchrony. Trends in Neurosciences, 24(10), 595–601. https://doi.org/10.1016/S0166-2236(00)01922-6

Jones, S. R., Pritchett, D. L., Sikora, M. A., Stufflebeam, S. M., Hämäläinen, M., & Moore, C. I. (2009). Quantitative Analysis and Biophysically Realistic Neural Modeling of the MEG Mu Rhythm: Rhythmogenesis and Modulation of Sensory-Evoked Responses. Journal of Neurophysiology, 102(6), 3554–3572. https://doi.org/10.1152/jn.00535.2009

Karvat, G., Schneider, A., Alyahyay, M., Steenbergen, F., Tangermann, M., & Diester, I. (2020). Real-time detection of neural oscillation bursts allows behaviourally relevant neurofeedback. Communications Biology, 3(1), Article 1. https://doi.org/10.1038/s42003-020-0801-z

Kehnemouyi, Y. M., Wilkins, K. B., Anidi, C. M., Anderson, R. W., Afzal, M. F., & Bronte-Stewart, H. M. (2021). Modulation of beta bursts in subthalamic sensorimotor circuits predicts improvement in bradykinesia. Brain, 144(2), 473–486. https://doi.org/10.1093/brain/awaa394

Keinrath, C., Wriessnegger, S., Müller-Putz, G. R., & Pfurtscheller, G. (2006). Postmovement beta synchronization after kinesthetic illusion, active and passive movements. International Journal of Psychophysiology, 62(2), 321–327. https://doi.org/10.1016/j.ijpsycho.2006.06.001

Khanna, P., & Carmena, J. M. (2017). Beta band oscillations in motor cortex reflect neural population signals that delay movement onset. ELife, 6, e24573. https://doi.org/10.7554/eLife.24573

Khawaldeh, S., Tinkhauser, G., Shah, S. A., Peterman, K., Debove, I., Nguyen, T. A. K., Nowacki, A., Lachenmayer, M. L., Schuepbach, M., Pollo, C., Krack, P., Woolrich, M., & Brown, P. (2020). Subthalamic nucleus activity dynamics and limb movement prediction in Parkinson’s disease. Brain, 143(2), 582–596. https://doi.org/10.1093/brain/awz417

Kilavik, B. E., Zaepffel, M., Brovelli, A., MacKay, W. A., & Riehle, A. (2013). The ups and downs of beta oscillations in sensorimotor cortex. Experimental Neurology, 245, 15–26. https://doi.org/10.1016/j.expneurol.2012.09.014

Kilner, J. M., Salenius, S., Baker, S. N., Jackson, A., Hari, R., & Lemon, R. N. (2003). Task-dependent modulations of cortical oscillatory activity in human subjects during a bimanual precision grip task. Neuroimage, 18(1), 67–73.

Kobak, D., Brendel, W., Constantinidis, C., Feierstein, C. E., Kepecs, A., Mainen, Z. F., Qi, X.-L., Romo, R., Uchida, N., & Machens, C. K. (2016). Demixed principal component analysis of neural population data. ELife, 5, e10989. https://doi.org/10.7554/eLife.10989

Kosciessa, J. Q., Grandy, T. H., Garrett, D. D., & Werkle-Bergner, M. (2020). Single-trial characterization of neural rhythms: Potential and challenges. NeuroImage, 206, 116331. https://doi.org/10.1016/j.neuroimage.2019.116331

Law, R. G., Pugliese, S., Shin, H., Sliva, D. D., Lee, S., Neymotin, S., Moore, C., & Jones, S. R. (2022). Thalamocortical Mechanisms Regulating the Relationship between Transient Beta Events and Human Tactile Perception. Cerebral Cortex, 32(4), 668–688. https://doi.org/10.1093/cercor/bhab221

Leocani, L., & Comi, G. (2006). Movement-related event-related desynchronization in neuropsychiatric disorders. In C. Neuper & W. Klimesch (Eds.), Progress in Brain Research (Vol. 159, pp. 351–366). Elsevier. https://doi.org/10.1016/S0079-6123(06)59023-5

Little, S., Bonaiuto, J., Barnes, G., & Bestmann, S. (2019). Human motor cortical beta bursts relate to movement planning and response errors. PLOS Biology, 17(10), e3000479. https://doi.org/10.1371/journal.pbio.3000479

Little, S., & Brown, P. (2014). The functional role of beta oscillations in Parkinson’s disease. Parkinsonism & Related Disorders, 20, S44–S48. https://doi.org/10.1016/S1353-8020(13)70013-0

Llera, A., Gómez, V., & Kappen, H. J. (2014). Adaptive multiclass classification for brain computer interfaces. Neural Computation, 26(6), 1108–1127. https://doi.org/10.1162/NECO_a_00592

Lofredi, R., Tan, H., Neumann, W.-J., Yeh, C.-H., Schneider, G.-H., Kühn, A. A., & Brown, P. (2019). Beta bursts during continuous movements accompany the velocity decrement in Parkinson’s disease patients. Neurobiology of Disease, 127, 462–471. https://doi.org/10.1016/j.nbd.2019.03.013

Lotte, F., Bougrain, L., Cichocki, A., Clerc, M., Congedo, M., Rakotomamonjy, A., & Yger, F. (2018). A review of classification algorithms for EEG-based brain-computer interfaces: A 10 year update. Journal of Neural Engineering, 15(3), 031005. https://doi.org/10.1088/1741-2552/aab2f2

Marshall, T. R., Quinn, A. J., Jensen, O., & Bergmann, T. O. (2022). Transcranial Direct Current Stimulation Alters the Waveform Shape of Cortical Gamma Oscillations [Preprint]. Neuroscience. https://doi.org/10.1101/2022.04.25.489371

McCarthy, M. M., Moore-Kochlacs, C., Gu, X., Boyden, E. S., Han, X., & Kopell, N. (2011). Striatal origin of the pathologic beta oscillations in Parkinson’s disease. Proceedings of the National Academy of Sciences, 108(28), 11620–11625. https://doi.org/10.1073/pnas.1107748108

McFarland, D. J., Miner, L. A., Vaughan, T. M., & Wolpaw, J. R. (2000). Mu and Beta Rhythm Topographies During Motor Imagery and Actual Movements. Brain Topography, 12(3), 177–186. https://doi.org/10.1023/A:1023437823106

Meirovitch, Y., Harris, H., Dayan, E., Arieli, A., & Flash, T. (2015). Alpha and Beta Band Event-Related Desynchronization Reflects Kinematic Regularities. Journal of Neuroscience, 35(4), 1627–1637. https://doi.org/10.1523/JNEUROSCI.5371-13.2015

Meyer, S. S., Bonaiuto, J., Lim, M., Rossiter, H., Waters, S., Bradbury, D., Bestmann, S., Brookes, M., Callaghan, M. F., Weiskopf, N., & Barnes, G. R. (2017). Flexible headcasts for high spatial precision MEG. Journal of Neuroscience Methods, 276, 38–45. https://doi.org/10.1016/j.jneumeth.2016.11.009

Miller, D. B., & O’Callaghan, J. P. (2015). Biomarkers of Parkinson’s disease: Present and future. Metabolism, 64(3, Supplement 1), S40–S46. https://doi.org/10.1016/j.metabol.2014.10.030

Miller, K. J., Schalk, G., Fetz, E. E., den Nijs, M., Ojemann, J. G., & Rao, R. P. N. (2010). Cortical activity during motor execution, motor imagery, and imagery-based online feedback. Proceedings of the National Academy of Sciences of the United States of America, 107(9), 4430–4435. https://doi.org/10.1073/pnas.0913697107

Mo, C., & Sherman, S. M. (2019). A Sensorimotor Pathway via Higher-Order Thalamus. Journal of Neuroscience, 39(4), 692–704. https://doi.org/10.1523/JNEUROSCI.1467-18.2018

Moca, V. V., Bârzan, H., Nagy-Dăbâcan, A., & Mureşan, R. C. (2021). Time-frequency super-resolution with superlets. Nature Communications, 12(1), 337. https://doi.org/10.1038/s41467-020-20539-9

Murthy, V. N., & Fetz, E. E. (1992). Coherent 25-to 35-Hz oscillations in the sensorimotor cortex of awake behaving monkeys. Proceedings of the National Academy of Sciences, 89(12), 5670–5674. https://doi.org/10.1073/pnas.89.12.5670

Nakagawa, K., Aokage, Y., Fukuri, T., Kawahara, Y., Hashizume, A., Kurisu, K., & Yuge, L. (2011). Neuromagnetic beta oscillation changes during motor imagery and motor execution of skilled movements: NeuroReport, 22(5), 217–222. https://doi.org/10.1097/WNR.0b013e328344b480

Neymotin, S. A., Daniels, D. S., Caldwell, B., McDougal, R. A., Carnevale, N. T., Jas, M., Moore, C. I., Hines, M. L., Hämäläinen, M., & Jones, S. R. (2020). Human Neocortical Neurosolver (HNN), a new software tool for interpreting the cellular and network origin of human MEG/EEG data. ELife, 9, e51214. https://doi.org/10.7554/eLife.51214

Papadopoulos, S., Bonaiuto, J., & Mattout, J. (2022). An Impending Paradigm Shift in Motor Imagery Based Brain-Computer Interfaces. Frontiers in Neuroscience, 15, 824759. https://doi.org/10.3389/fnins.2021.824759

Pauls, K. A. M., Korsun, O., Nenonen, J., Nurminen, J., Liljeström, M., Kujala, J., Pekkonen, E., & Renvall, H. (2022). Cortical beta burst dynamics are altered in Parkinson’s disease but normalized by deep brain stimulation. NeuroImage, 257, 119308. https://doi.org/10.1016/j.neuroimage.2022.119308

Pedregosa, F., Varoquaux, G., Gramfort, A., Michel, V., Thirion, B., Grisel, O., Blondel, M., Prettenhofer, P., Weiss, R., & Dubourg, V. (2011). Scikit-learn: Machine learning in Python. Journal of Machine Learning Research, 12(Oct), 2825–2830.

Pelli, D. G. (1997). The VideoToolbox software for visual psychophysics: Transforming numbers into movies. Spatial Vision.

Perone, S., & Gartstein, M. A. (2019). Mapping cortical rhythms to infant behavioral tendencies via baseline EEG and parent-report. Developmental Psychobiology, 61(6), 815–823. https://doi.org/10.1002/dev.21867

Pfurtscheller, G. (1981). Central beta rhythm during sensorimotor activities in man. Electroencephalography and Clinical Neurophysiology, 51(3), 253–264. https://doi.org/10.1016/0013-4694(81)90139-5

Pfurtscheller, G., & Lopes da Silva, F. H. (1999). Event-related EEG/MEG synchronization and desynchronization: Basic principles. Clinical Neurophysiology, 110(11), 1842–1857. https://doi.org/10.1016/S1388-2457(99)00141-8

Pfurtscheller, G., & Neuper, C. (2001). Motor imagery and direct brain-computer communication. Proceedings of the IEEE, 89(7), 1123–1134. https://doi.org/10.1109/5.939829

Pfurtscheller, G., Stancák, A., & Edlinger, G. (1997). On the existence of different types of central beta rhythms below 30 Hz. Electroencephalography and Clinical Neurophysiology, 102(4), 316–325. https://doi.org/10.1016/S0013-4694(96)96612-2

Pfurtscheller, G., Stancák, A., & Neuper, Ch. (1996). Event-related synchronization (ERS) in the alpha band — an electrophysiological correlate of cortical idling: A review. International Journal of Psychophysiology, 24(1), 39–46. https://doi.org/10.1016/S0167-8760(96)00066-9

Picazio, S., Veniero, D., Ponzo, V., Caltagirone, C., Gross, J., Thut, G., & Koch, G. (2014). Prefrontal Control over Motor Cortex Cycles at Beta Frequency during Movement Inhibition. Current Biology, 24(24), 2940–2945. https://doi.org/10.1016/j.cub.2014.10.043

Pogosyan, A., Gaynor, L. D., Eusebio, A., & Brown, P. (2009). Boosting Cortical Activity at Beta-Band Frequencies Slows Movement in Humans. Current Biology, 19(19), 1637–1641. https://doi.org/10.1016/j.cub.2009.07.074

Pollok, B., Latz, D., Krause, V., Butz, M., & Schnitzler, A. (2014). Changes of motor-cortical oscillations associated with motor learning. Neuroscience, 275, 47–53. https://doi.org/10.1016/j.neuroscience.2014.06.008

Porr, B., & Howell, L. (2019). R-peak detector stress test with a new noisy ECG database reveals significant performance differences amongst popular detectors [Preprint]. Bioengineering. https://doi.org/10.1101/722397

Pratt, J. W., & Gibbons, J. D. (1981). Kolmogorov-Smirnov Two-Sample Tests. In J. W. Pratt & J. D. Gibbons, Concepts of Nonparametric Theory (pp. 318–344). Springer New York.

Quinn, A. J., Lopes-dos-Santos, V., Huang, N., Liang, W.-K., Juan, C.-H., Yeh, J.-R., Nobre, A. C., Dupret, D., & Woolrich, M. W. (2021). Within-cycle instantaneous frequency profiles report oscillatory waveform dynamics. BioRxiv, 2021.04.12.439547. https://doi.org/10.1101/2021.04.12.439547

R Core Team. (2022). R: A language and environment for statistical computing. R Foundation for Statistical Computing.

Rayson, H., Debnath, R., Alavizadeh, S., Fox, N., Ferrari, P. F., & Bonaiuto, J. J. (2022). Detection and analysis of cortical beta bursts in developmental EEG data. Developmental Cognitive Neuroscience, 54, 101069. https://doi.org/10.1016/j.dcn.2022.101069

Rempe, M. P., Lew, B. J., Embury, C. M., Christopher-Hayes, N. J., Schantell, M., & Wilson, T. W. (2022). Spontaneous sensorimotor beta power and cortical thickness uniquely predict motor function in healthy aging. NeuroImage, 263, 119651. https://doi.org/10.1016/j.neuroimage.2022.119651

Reuter, E.-M., Booms, A., & Leow, L.-A. (2022). Using EEG to study sensorimotor adaptation. Neuroscience & Biobehavioral Reviews, 134, 104520. https://doi.org/10.1016/j.neubiorev.2021.104520

Rhodes, E., Gaetz, W. C., Marsden, J., & Hall, S. D. (2018). Transient Alpha and Beta Synchrony Underlies Preparatory Recruitment of Directional Motor Networks. Journal of Cognitive Neuroscience, 30(6), 867–875. https://doi.org/10.1162/jocn_a_01250

Ridgway, G. R., Litvak, V., Flandin, G., Friston, K. J., & Penny, W. D. (2012). The problem of low variance voxels in statistical parametric mapping; a new hat avoids a ‘haircut’. NeuroImage, 59(3), 2131–2141. https://doi.org/10.1016/j.neuroimage.2011.10.027

Rossi, C., & Van Schependom, J. (2022). Two approaches to tackle the sign ambiguity of beamformed MEG source-reconstructed data. International Conference on Biomagnetism.

Rossiter, H. E., Davis, E. M., Clark, E. V., Boudrias, M.-H., & Ward, N. S. (2014). Beta oscillations reflect changes in motor cortex inhibition in healthy ageing. NeuroImage, 91, 360–365. https://doi.org/10.1016/j.neuroimage.2014.01.012

Salenius, S., & Hari, R. (2003). Synchronous cortical oscillatory activity during motor action. Current Opinion in Neurobiology, 13(6), 678–684. https://doi.org/10.1016/j.conb.2003.10.008

Schmidt, S. L., Peters, J. J., Turner, D. A., & Grill, W. M. (2020). Continuous deep brain stimulation of the subthalamic nucleus may not modulate beta bursts in patients with Parkinson’s disease. Brain Stimulation, 13(2), 433–443. https://doi.org/10.1016/j.brs.2019.12.008

Sherman, M. A., Lee, S., Law, R., Haegens, S., Thorn, C. A., Hämäläinen, M. S., Moore, C. I., & Jones, S. R. (2016). Neural mechanisms of transient neocortical beta rhythms: Converging evidence from humans, computational modeling, monkeys, and mice. Proceedings of the National Academy of Sciences, 113(33), E4885–E4894. https://doi.org/10.1073/pnas.1604135113

Shin, H., Law, R., Tsutsui, S., Moore, C. I., & Jones, S. R. (2017). The rate of transient beta frequency events predicts behavior across tasks and species. ELife, 6, e29086. https://doi.org/10.7554/eLife.29086

Smith, S. M., & Nichols, T. E. (2009). Threshold-free cluster enhancement: Addressing problems of smoothing, threshold dependence and localisation in cluster inference. NeuroImage, 44(1), 83–98. https://doi.org/10.1016/j.neuroimage.2008.03.061

Song, X., Yoon, S.-C., & Perera, V. (2013). Adaptive Common Spatial Pattern for single-trial EEG classification in multisubject BCI. 2013 6th International IEEE/EMBS Conference on Neural Engineering (NER), 411–414. https://doi.org/10.1109/NER.2013.6695959

Sporn, S., Hein, T., & Herrojo Ruiz, M. (2020). Alterations in the amplitude and burst rate of beta oscillations impair reward-dependent motor learning in anxiety. ELife, 9, e50654. https://doi.org/10.7554/eLife.50654

Tan, H., Jenkinson, N., & Brown, P. (2014). Dynamic neural correlates of motor error monitoring and adaptation during trial-to-trial learning. The Journal of Neuroscience: The Official Journal of the Society for Neuroscience, 34(16), 5678–5688. https://doi.org/10.1523/JNEUROSCI.4739-13.2014

Tan, H., Wade, C., & Brown, P. (2016). Post-Movement Beta Activity in Sensorimotor Cortex Indexes Confidence in the Estimations from Internal Models. Journal of Neuroscience, 36(5), 1516–1528. https://doi.org/10.1523/JNEUROSCI.3204-15.2016

Thaler, L., Schütz, A. C., Goodale, M. A., & Gegenfurtner, K. R. (2013). What is the best fixation target? The effect of target shape on stability of fixational eye movements. Vision Research, 76, 31–42. https://doi.org/10.1016/j.visres.2012.10.012

Tinkhauser, G., Pogosyan, A., Little, S., Beudel, M., Herz, D. M., Tan, H., & Brown, P. (2017). The modulatory effect of adaptive deep brain stimulation on beta bursts in Parkinson’s disease. Brain: A Journal of Neurology, 140(4), 1053–1067. https://doi.org/10.1093/brain/awx010

Tinkhauser, G., Pogosyan, A., Tan, H., Herz, D. M., Kühn, A. A., & Brown, P. (2017). Beta burst dynamics in Parkinson’s disease OFF and ON dopaminergic medication. Brain, 140(11), 2968–2981. https://doi.org/10.1093/brain/awx252

Titova, N., & Chaudhuri, K. R. (2017). Personalized medicine in Parkinson’s disease: Time to be precise. Movement Disorders, 32(8), 1147–1154. https://doi.org/10.1002/mds.27027

Torrecillos, F., Tinkhauser, G., Fischer, P., Green, A. L., Aziz, T. Z., Foltynie, T., Limousin, P., Zrinzo, L., Ashkan, K., Brown, P., & Tan, H. (2018). Modulation of Beta Bursts in the Subthalamic Nucleus Predicts Motor Performance. Journal of Neuroscience, 38(41), 8905–8917. https://doi.org/10.1523/JNEUROSCI.1314-18.2018

Trevarrow, M. P., Kurz, M. J., McDermott, T. J., Wiesman, A. I., Mills, M. S., Wang, Y.-P., Calhoun, V. D., Stephen, J. M., & Wilson, T. W. (2019). The developmental trajectory of sensorimotor cortical oscillations. NeuroImage, 184, 455–461. https://doi.org/10.1016/j.neuroimage.2018.09.018

Tzagarakis, C., Ince, N. F., Leuthold, A. C., & Pellizzer, G. (2010). Beta-Band Activity during Motor Planning Reflects Response Uncertainty. Journal of Neuroscience, 30(34), 11270–11277. https://doi.org/10.1523/JNEUROSCI.6026-09.2010

Tzagarakis, C., West, S., & Pellizzer, G. (2015). Brain oscillatory activity during motor preparation: Effect of directional uncertainty on beta, but not alpha, frequency band. Frontiers in Neuroscience, 9. https://doi.org/10.3389/fnins.2015.00246

van Wijk, B. C. M., Daffertshofer, A., Roach, N., & Praamstra, P. (2009). A Role of Beta Oscillatory Synchrony in Biasing Response Competition? Cerebral Cortex, 19(6), 1294–1302. https://doi.org/10.1093/cercor/bhn174

Vidaurre, D., Quinn, A. J., Baker, A. P., Dupret, D., Tejero-Cantero, A., & Woolrich, M. W. (2016). Spectrally resolved fast transient brain states in electrophysiological data. NeuroImage, 126, 81–95. https://doi.org/10.1016/j.neuroimage.2015.11.047

Vieira, V. M. N. C. S. (2012). Permutation tests to estimate significances on Principal Components Analysis. Computational Ecology and Software, 2(2), 103–124.

Walsh, C., Ridler, T., Margetts-Smith, G., Garrido, M. G., Witton, J., Randall, A. D., & Brown, J. T. (2022). Beta bursting in the retrosplenial cortex is a neurophysiological correlate of environmental novelty which is disrupted in a mouse model of Alzheimer’s disease. Journal of Neuroscience. https://doi.org/10.1523/JNEUROSCI.0890-21.2022

Watson, A. B., & Pelli, D. G. (1983). Quest: A Bayesian adaptive psychometric method. Perception & Psychophysics, 33(2), 113–120. https://doi.org/10.3758/BF03202828

Wessel, J. R. (2020). β-Bursts Reveal the Trial-to-Trial Dynamics of Movement Initiation and Cancellation. Journal of Neuroscience, 40(2), 411–423. https://doi.org/10.1523/JNEUROSCI.1887-19.2019

Wessel, J. R., & Aron, A. R. (2017). On the Globality of Motor Suppression: Unexpected Events and Their Influence on Behavior and Cognition. Neuron, 93(2), 259–280. https://doi.org/10.1016/j.neuron.2016.12.013

West, T. O., Duchet, B., Farmer, S. F., Friston, K. J., & Cagnan, H. (2022). When do bursts matter in the motor cortex? Investigating changes in the intermittencies of beta rhythms associated with movement states [Preprint]. Neuroscience. https://doi.org/10.1101/2022.06.22.497199

Wu, S., & Wang, J. (2014). Nonnegative matrix factorization: When data is not nonnegative. 2014 7th International Conference on Biomedical Engineering and Informatics, 227–231. https://doi.org/10.1109/BMEI.2014.7002775

Yeh, C.-H., Al-Fatly, B., Kühn, A. A., Meidahl, A. C., Tinkhauser, G., Tan, H., & Brown, P. (2020). Waveform changes with the evolution of beta bursts in the human subthalamic nucleus. Clinical Neurophysiology, 131(9), 2086–2099. https://doi.org/10.1016/j.clinph.2020.05.035

Zhang, W., & Bruno, R. M. (2019). High-order thalamic inputs to primary somatosensory cortex are stronger and longer lasting than cortical inputs. ELife, 8, e44158. https://doi.org/10.7554/eLife.44158

Zhang, Y., Chen, Y., Bressler, S. L., & Ding, M. (2008). Response preparation and inhibition: The role of the cortical sensorimotor beta rhythm. Neuroscience, 156(1), 238–246. https://doi.org/10.1016/j.neuroscience.2008.06.061

Zich, C., Quinn, A. J., Bonaiuto, J. J., O’Neill, G., Mardell, L. C., Ward, N. S., & Bestmann, S. (2022). Spatiotemporal organization of human sensorimotor beta burst activity. BioRxiv.

Zich, C., Quinn, A. J., Mardell, L. C., Ward, N. S., & Bestmann, S. (2020). Dissecting Transient Burst Events. Trends in Cognitive Sciences, 24(10), 784–788. https://doi.org/10.1016/j.tics.2020.07.004

